# Type 1 immunity enables neonatal thymic ILC1 production

**DOI:** 10.1101/2023.02.28.530451

**Authors:** Peter Tougaard, Mario Ruiz Pérez, Wolf Steels, Jelle Huysentruyt, Bruno Verstraeten, Jessica Vetters, Tatyana Divert, Amanda Gonçalves, Ria Roelandt, Nozomi Takahashi, Sophie Janssens, Terkild Brink Buus, Tom Taghon, Georges Leclercq, Peter Vandenabeele

## Abstract

Thymic atrophy occurs following type 1 inflammatory conditions like viral infection and sepsis, resulting in cell death and disruption of T-cell development. However, it remains undetermined whether the thymus actively contributes to the immune response. Thus, we cultured neonatal thymus *ex vivo* with the type 1 cytokines IL-12 plus IL-18, resulting in a rapid shift from steady-state T-cell development to the production, expansion, and thymic exit of CXCR6^+^CD62L^-^ type 1 innate lymphoid cells (ILC1s). Single-cell RNA-sequencing and functional assays identified these cells as embryonic-wave-derived KLRG1^+^ ILC1s that mainly differentiated from immature neonatal thymic ILC1s. Confocal 3D imaging confirmed neonatal thymic ILC1 expansion during MCMV infection. Furthermore, thymic grafts revealed *in vivo* thymic ILC1 egress and type 1 inflammation-induced homing of thymus-derived KLRG1^+^ ILC1s to the liver and peritoneal cavity. Altogether, our data reveal a novel thymic function where type 1 immunity enables the production and peripheral homing of thymic-derived ILC1s.

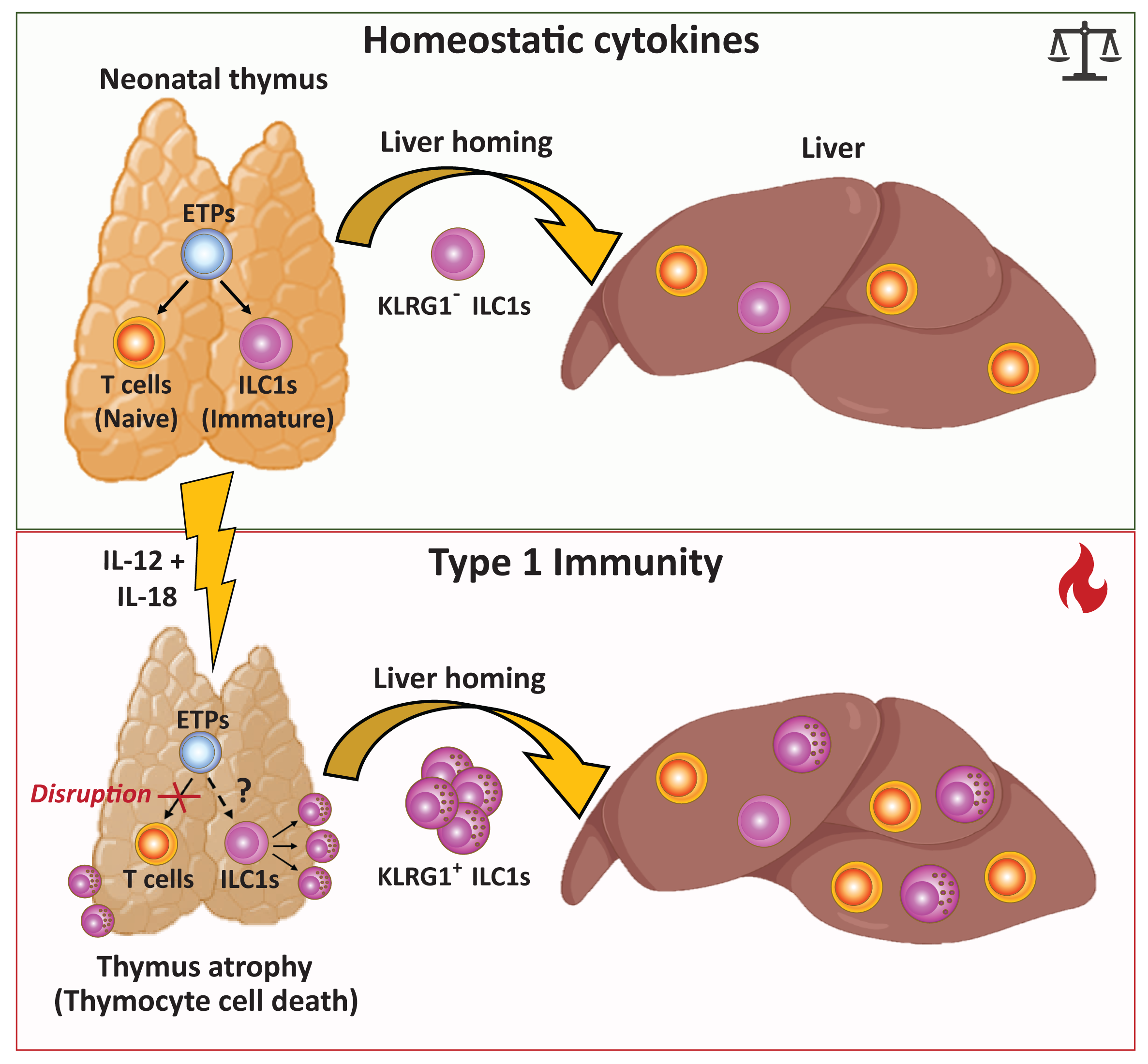

---

The type 1 immunity-inducing cytokines IL-12 and IL-18 are critical in viral infection^1, 2^ and sepsis^3, 4^. Human and murine cytomegalovirus (CMV) infections are associated with increased production of IL-12 and IL-18^2, 5, 6^, while congenital and perinatal human CMV infections can result in morbidity or life-long neurological complications^7, 8^. Although neonatal CMV infection rarely results in long-term complications, preterm infants can develop serious sepsis-like syndrome^8, 9^. During viral infection and sepsis, the thymus undergoes acute thymic atrophy^10, 11^, where T cell development is disrupted but can recover when the inflammation is resolved^12, 13^. Acute thymic atrophy differs from age-related thymic involution, where the thymic tissue is slowly replaced with adipose tissue^14^. Although infection-induced thymic atrophy is an evolutionarily conserved process^11, 15^, the underlying causality and selective advantage for the host have not been determined.

Group-1 innate lymphoid cells (ILCs) comprise ILC1s and conventional NK (cNK) cells. ILC1s primarily reside in the tissues and mediate an early response against viral infections and cancer, whereas cNK cells circulate and act throughout the immune response^16–18^. ILC1s are seeded in the body during several perinatal waves^19, 20^ and, in mice, they constitute the main population of group-1 ILCs across tissues at birth^19^. Nevertheless, the ontogeny of the different peripheral ILC1 waves remains elusive^19, 21^. cNK cells originate from the bone marrow and develop postnatally to become the dominant group-1 ILC population in the spleen and bone marrow within the first two weeks of life^19^. Thymic group-1 ILCs are recognized as a rare population, distinct from other peripheral group-1 ILCs^22, 23^, and can develop from early thymic progenitors (ETPs)^24–27^. Unlike cNK cells, thymic group-1 ILC development depends on IL-7 and the transcription factor GATA3^22^, similar to ILC1s, ILC2s, and ILC3s^28^. ILC1 numbers increase in all examined tissues during perinatal development in mice from embryonic day 14.5 (E14.5) until postnatal day 3 (P3)^19^. The only exception is the thymus, where ILC1 cell numbers reach maximum levels on the day of birth and decrease between P1 and P3^19^. Whether this thymic-specific decrease in postnatal ILC1s signifies cell death or egress to peripheral tissues remains to be determined.

ILC1s and cNK cells express classical pan-NK cell markers, such as NKp46, CD122, and NK1.1, while being negatively delineated for T-and B-cell lineage markers^19, 28, 29^. Most ILC1s are CD49a^+^CD49b^+^ double positive at birth, with CD49b expression decreasing gradually in the first weeks of life^19^. During viral infection and in cancer models in adult mice, cNK cells acquire “ILC1-like” features, such as tissue residency and enhanced cytokine production^30–32^. Despite their phenotypical overlap, ILC1s and cNK cells are considered separate lineages with no developmental interconversion^19, 29, 33–35^. ILC1s that display a high capacity for cytokine production and low levels of granzyme have been coined “helper-like” ILC1s based on the parallels with CD4 helper T cells^29, 36^. Moreover, IL-7R^+^ helper-like ILC1s represent a less mature state of ILC1s that can differentiate into Granzyme-B^+^IL-7R^-^ “cytotoxic” ILC1s (Friedrich *et al., Nature Immunology* (2021)). This observation challenges the classical delineation between ILC1s and cNK cells based on their respective capacity for cytokine production and cytotoxic killing^19, 36^.

Here, we investigated the effect of acute inflammation on the neonatal thymus by *in vivo* murine cytomegalovirus (MCMV) infection and during sterile inflammation by administering combined IL-12 plus IL-18. We document that MCMV infection and IL-12+IL-18 injections both result in acute thymic atrophy in neonatal mice. Using neonatal thymus organ cultures (NTOCs), we show that the combined administration of IL-12+IL-18 results in the expansion of thymic-exiting KLRG1^+^ ILC1s that primarily differentiates from immature thymic ILC1s. Fate-mapping *Ncr1*-tdTomato mice were used to visualize the *in situ* expansion of these ILC1s in the thymus of MCMV-infected neonates, and subcapsular kidney grafting shows that thymic ILC1s can egress the thymus and upon *in vivo* administration of type 1 cytokines expand rapidly and home to the liver and peritoneal cavity. Altogether, our results demonstrate a novel thymic function where type 1 immunity enables the thymus to rapidly produce peripheral homing ILC1s. Thereby, revealing that the thymus can produce other peripheral homing lymphocytes than T cells.

## Results

### Type-1 inflammation induces thymus-exiting ILC1s during NTOC

Infecting neonatal mice with MCMV results in acute thymus atrophy within five days **(Figures 1A-1B)**. MCMV infection rapidly induces high levels of the type-1 cytokines IL-12 and IL-18^2^. Since acute thymic atrophy is associated with conditions with enhanced cytokine responses^10, 11, 37^, we investigated whether the injection of IL-12 and IL-18 could cause thymic atrophy during sterile inflammation. Similarly to MCMV infection (**Figures 1A-1B**), i.p. injection of IL-12 and IL-18 in both neonatal and adult mice for three consecutive days resulted in severe acute thymus atrophy, here shown on Day 5 **(Figures 1C-1D and S1A-S1B)**. To investigate the direct effect of IL-12+IL-18 on the thymus, we cultured neonatal thymic lobes in a neonatal thymic organ culture (NTOC) in the presence of the two cytokines (**Figure 1E**). Similar to what was observed *in vivo*, IL-12+IL-18 administration resulted in cell loss as compared to the vehicle condition (**Figures 1F** **and S1C)**. Surprisingly, we observed a large population of group-1 ILCs (either ILC1s or cNK cells) exiting into the supernatant of the organ culture within six days and comprising approximately 25% of the thymus-exiting cells. Furthermore, various myeloid subsets were also observed exiting from the thymus into the supernatant in all four conditions (**Figures 1F** **and S1C-D)**. The group-1 ILCs expressed CD122^+^IL-18Rα^+^NK-1.1^+^ and were lineage-negative (Lin-: TCRβ^-^TCRγδ^-^CD3e^-^CD4^-^CD8β^-^Ly-6G^-^CD19^-^ CD115^-^CD11c^-^) (**Figures 1F** **and S1C)**. As ILC1s express CD49b at birth, it is not an exclusive cNK cell marker^19^, but ILC1s are CD49a^+^CD62L^-^ ^19, 21, 28^. Thus, when comparing the NTOC-exiting group-1 ILCs with eight-week-old adults and P0.5 neonatal thymic group-1 ILCs (CD122^+^NK1.1^+^Lin^-^ cells), we confirmed them as ILC1s based on their CD49a^+^CD62L^-^ phenotype. The neonatal thymus ILC1s had a similar CD49a^+^CD62L^-^ phenotype as the NTOC-derived ILC1 population. Only a minority of the adult thymic group-1 ILCs had the CD49a^+^CD62L^-^ ILC1 phenotype observed in neonatal mice (**Figure 1G**). Although Eomes and CD49a are co-expressed in some ILC1 subsets, like the salivary gland ILC1s, Eomes and CD49a expression are mutually exclusive for hepatic cNKs and ILC1s, respectively^21, 36^. Thus to uniformly identify ILC1s and cNK cells across tissues in neonates and adult mice, we gated on CD122^+^NK1.1^+^Lin^-^ cells and defined ILC1s as (CD49a^+^CD62L^-^) and cNK cells as (CD49b^+^CD49a^-^), verifyng this definition based on intracellular Eomes expression **(Figures S1E and S1F)**. Apart from ILC1s and cNK cells, a subset of NK cells in the adult thymus could be defined as (CD49b^+^CD49a^+^CD62L^+^) and displayed high Eomes expression **(Figures S1E and S1F)**, as previously decribed^23^. Contrary to adult thymus ILC1s, both the neonatal thymus ILC1s and NTOC-derived ILC1s had a low expression of Eomes **(Figure S1E)**. Kinetic measurements revealed that most of the initial exit and expansion of thymic-derived ILC1s in the supernatant of IL-12+IL-18-treated NTOC occurred rapidly in the first three days after the organ culture was set up (**Figure 1H**). Electron microscopy of sorted ILC1s and T cells from NTOC supernatant revealed a distinct morphology of the thymic-exiting ILC1s (**Figure 1I**). After six days, protein measurement of the cytokines secreted in the NTOC supernatant revealed high levels of IFN-γ, TNF-α, CCL3, and GM-CSF following IL-12+IL-18 stimulation **(Figure S1G)**. We performed intracellular flow cytometry on IL-12+IL-18-treated NTOC supernatant cells showed enhanced production of IFN-γ, TNF-α, GM-CSF and Granzyme B in the ILC1 population, verifying them as cytokine-producing **(Figure S1H)**. To further determine if the thymus-exiting ILC1s had cytotoxic capacity, we sorted ILC1s from IL-12+IL-18-treated NTOC. NTOC ILC1s exhibited a dose-dependent killing of YAC-1 target cells, indicating these cells to be cytotoxic ILC1s (**Figure 1J**). Finally, to further investigate the NTOC-exiting ILC1s, cellular indexing of transcriptomes and epitopes by sequencing (CITE-seq) analysis was performed on viable cells from the supernatant following six days of NTOC (**Figure 1K**). Twenty-three cell populations were identified from the four conditions of NTOC-exiting cells (Vehicle, IL-12, IL-18, and IL-12+IL-18) **(Figures S1I-S1K)**. Based on the top log-fold differentially expressed (DE)-genes and manually curated genes **(Figures S1I and S1J)**, we identified the overall cell type of each of the thymus-derived populations (**Figures 1L** **and S1K).** The lack of *Cd3e*, *Trdc*, *Trac* and *Bcl11b* expression further confirmed the expanded NTOC population as ILC1s rather than T cells. The expression of *Ncr1*, *Tbx21*, *Cxcr6*, *Gzmb, Il2rb*, and the absence of *Sell* (encoding CD62L) corroborated their ILC1 identity along with their cytotoxic phenotype **(Figure S1J)**. The single-cell data showed a large population of thymic-derived ILC1s exiting the NTOC in response to IL-12+IL-18 stimulation (**Figure 1M**). Along with the ILC1s, several large populations of granulocytes, monocytes and macrophages were also observed to exit the NTOC lobes (**Figure 1L**), confirming findings by flow cytometry **(Figures 1M, S1I, and S1J)**. In addition, CITE-seq antibody labeling revealed the group-1 ILC differentiation marker KLRG1^38^ to be highly expressed on the thymic-exiting ILC1s following IL-12+IL-18 stimulation **(Figure S1L)**.

**Figure 1:**
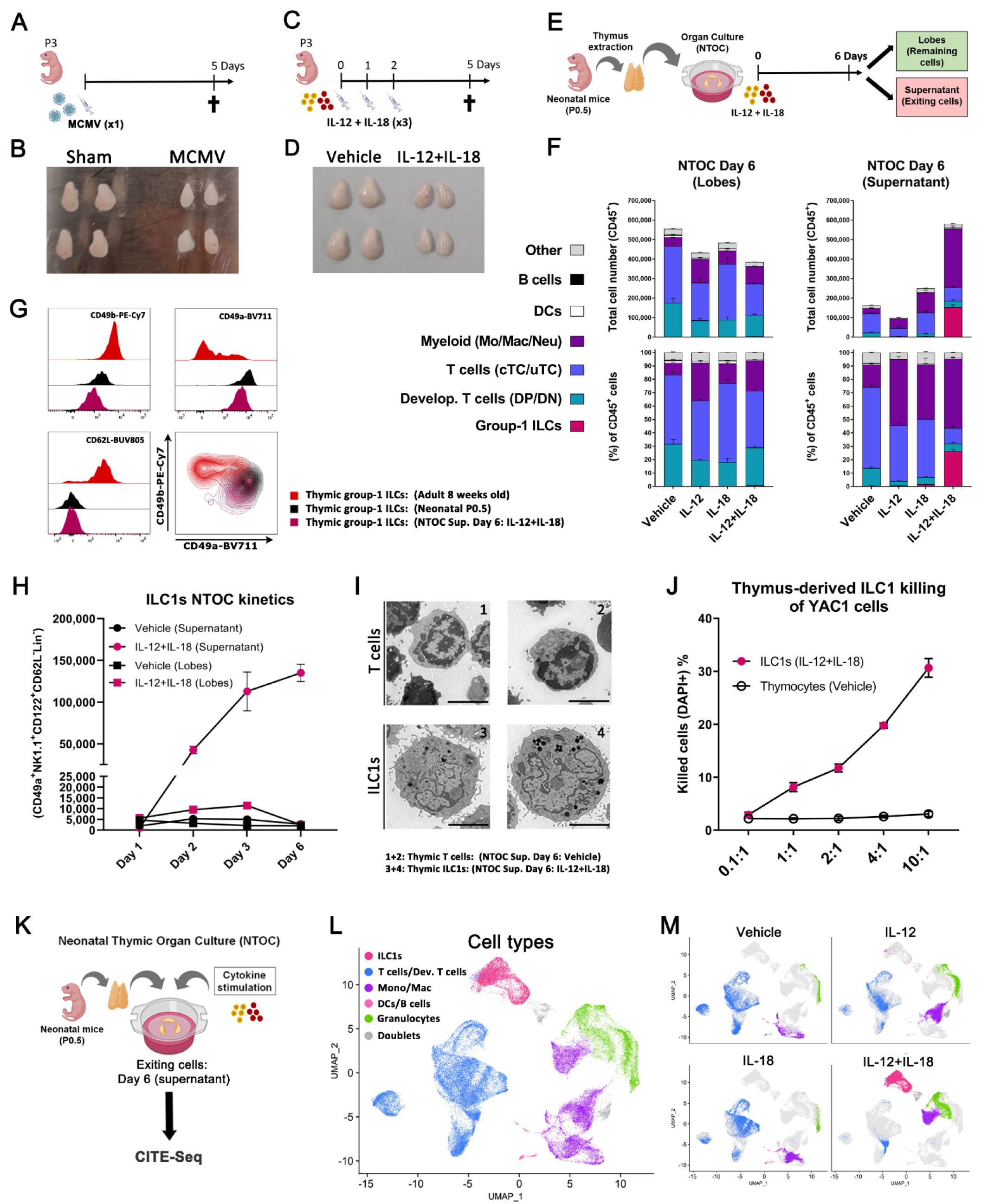
Type-1 inflammation induces expansion of thymic-exiting ILC1s from NTOC. **(A)** Schematic of mice infected once on perinatal day 3 (P3) with 3000 pfu MCMV and terminated after 5 days. **(B)** As shown in **A**, pictures of thymic lobes 5 days after infection with MCMV or Sham are representative of four independent experiments. **(C)** Schematic of mice injected three times from P3 to P5 with IL12+IL-18 and terminated 5 days after the first injection. **(D)** As shown in **C**, a picture of both thymic lobes from two neonates 5 days after the first injection with IL-12+IL-18 or Vehicle representative of three independent experiments. **(E)** Schematic of NTOC-strategy used to analyze cells in **F**, using intact neonatal thymus lobes cultured on a floating filter and stimulated with cytokines followed by analyzing the cells inside the lobes and the cells that exited into the supernatant. **(F)** Stacked bar plot of neonatal thymic organ culture (NTOC) performed as illustrated in E, showing the total number (upper) and percentage (lower) of CD45^+^ cell types inside the thymic lobes (left) and the cells that have exited the lobes into the supernatant (right). **(G)** Flow cytometry comparison of using ILC1 and cNK cell markers on group-1 ILCs (CD122^+^NK1.1^+^Lin^-^) from the thymus in adults (8-weeks-old), neonates (P0.5), and IL-12+IL-18-stimulated NTOC supernatant (Day 6). Thymus-derived group-1 ILCs are shown as histograms and density plots (showing outliers down to the 2^nd^ percentile). **(H)** Kinetic overview of the ILC1 expansion in the NTOC inside the lobes and in the supernatant, comparing Vehicle control with IL-12+IL-18 stimulation from NTOC Days 1, 2, 3, and 6. **(I)** Representative transmission electron microscopy (TEM) pictures of cell sorted NTOC-derived CD4 T cells (1 and 2; Vehicle supernatant) or ILC1s (3 and 4; IL-12+IL-18 supernatant). The bars in the bottom right corner are 2 µm. **(J)** YAC1 killing assay, showing dose-dependent killing with increasing cell numbers of sorted effector cells (ILC1s or control thymocytes) and constant amount of target cells (YAC1 cells). Killing was shown as DAPI^+^ YAC1 cells after 4 hours of co-culture. **(K)** Schematic of Day 6 NTOC supernatant cells used for CITE-seq. DAPI-cells were sorted from the supernatant from four conditions: **1)** Vehicle, **2)** IL-18, **3)** IL-12, **4)** IL-12+IL-18. **(L)** UMAP plot of scRNAseq data, dividing the cells into major cell types observed. **(M)** Split UMAPs of the four conditions applying the same cell type colors as the overview in **L**. **(F and H)** Error bars in results are shown as SEM from three biological replicates (*n*=3) and representative of at least five independent experiments. **(G)** Results representative of six biological replicates from two independent experiments. **(J)** Results are shown as SEM from three technical replicates (*n*=3) and representative of three independent experiments. **Abbreviations:** CITE-seq = Cellular Indexing of Transcriptomes and Epitopes by Sequencing. Lin^-^ = Lineage negative, defined as **(F)** TCRβ^-^TCRγδ^-^CD3e^-^CD4^-^CD8β^-^Ter-119^-^Ly-6G^-^CD19^-^CD11c^-^CD115^-^, and **(G and H)** TCRβ^-^TCRγδ^-^CD3e^-^CD4^-^CD8β^-^Ter-119^-^Ly-6G^-^CD19^-^CD11c^-^F4/80^-^. MCMV = Murine cytomegalovirus, pfu = Plague forming units. NTOC = neonatal thymic organ culture. UMAP = Uniform Manifold Approximation and Projection.

### KLRG1^+^ ILC1s differentiate and expand from immature thymic ILC1s

Neonatal thymus lobes from *Rag2*^-/-^OTI TCR-transgene mice still showed NTOC expansion of ILC1s **(Figure S2A)**, thereby confirming the ontogenesis of thymic-derived ILC1s being independent from T cell development^24^. Furthermore, we confirmed high expression of the differentiation marker KLRG1^38^ (KLRG1^hi^) following IL-12+IL-18 stimulation, whereas vehicle control and *ex vivo* extracted ILC1s display a more immature CD27^+/low^KLRG1^-^ phenotype (**Figure 2A**). Thymus seeding ETPs have multipotent potential^39^. However, it has not been clarified whether ETP-derived group-1 ILCs are ILC1s or cNK cells. Thus, we hypothesize that the KLRG1^hi^ ILC1s either differentiate directly from ETPs, uncommitted double negative 2a (DN2a) cells, or from immature thymic ILC1s (**Figure 2B**). Thus, we sorted ETPs (CD117^hi^CD44^+^CD25^-^Lin^-^), DN2a (CD117^hi^CD44^+^CD25^+^Lin^-^) and the rare ILC1s (CD122^+^Lin^-^) from *ex vivo* neonatal (P0.5) thymus. From each of these sorted cell populations, 1000 cells were co-cultured on OP9-DL1 cells with different combinations of the common γ-chain cytokines (IL-2, IL-7, and IL-15) in the presence or absence of IL-12+IL-18 to reveal the precursor of the KLRG1^hi^ expanding ILC1s (**Figure 2C**). We found that ETPs could develop into group-1 ILCs in a highly IL-7-dependent manner (**Figure 2D**), confirming previous findings^22, 24, 26^. Their ILC1 identity was confirmed by their CD122^+^CD49a^+^CD62L^-^ phenotype, and neonatal DN2a seeded cells consistently only produced few ILC1s (**Figures 2D** **and S2B)**, indicating a notable loss of ILC1-potential compared to ETPs. ETP-derived ILC1s primarily displayed a KLRG1^-^CD27^+^ phenotype, both in the presence and absence of IL-12+IL-18 (**Figures 2D** **and 2E)**. Finally, seeding neonatal thymic ILC1s followed by six days of OP9-DL1 culture with IL-2+IL-7+IL-15 mainly yielded an expansion of the immature KLRG1^-^CD27^+^ ILC1s. However, the addition of IL-12+IL-18 primarily resulted in the differentiation to KLRG1^+^CD27^-^ ILC1s (**Figure 2D, third panel and** **Figure 2E**). These results indicate that most of the observed NTOC-exiting KLRG1^hi^ ILC1s are likely differentiating and expanding from immature KLRG1^-^ thymic ILC1s rather than ETPs.

**Figure 2:**
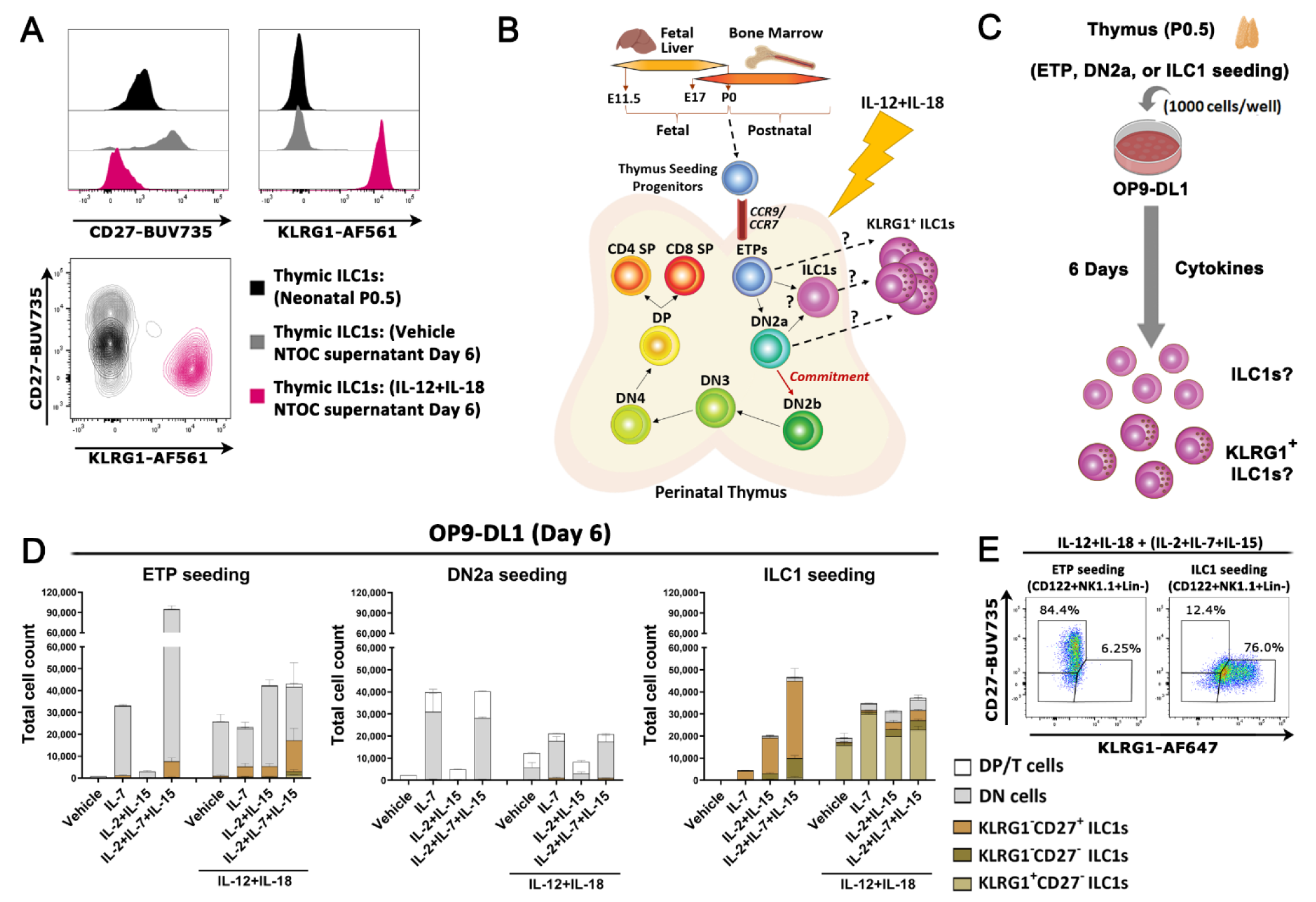
IL-12+IL-18-induced thymic ILC1s expand from immature ILC1s. **(A)** Flow cytometry comparison of neonatal thymic ILC1s from Day 4 NTOC or *ex vivo* P0.5 and their expression of CD27 and KLRG1. Data are either shown as histograms or density plots (showing outliers down to the 2^nd^ percentile). **(B)** Schematic of thymic development and T cell commitment at the DN2b stage, visualizing hypothesized precursors for the KLRG1^+^ ILC1s. **(C)** Schematic of OP9-DL1 organ culture setup to test the hypothesis illustrated in **B**, where 1000 cell sorted progenitor cells were added to each 96-well on top of OP9-DL1 cells to determine whether the rapidly expanded KLRG1^+^ ILC1s derived from neonatal (P0.5) early thymic progenitors (ETPs) or immature ILC1s. DL1 notch-ligand-expressing OP9 cells were used to simulate T cell promoting thymic environment, to make sure T cell development remained possible. **(D)** Stacked bar plots showing developing T cells or ILC1s based on their expression of KLRG1 and CD27 following 6 days of OP9-DL1 co-culture for ETPs (CD117^(hi)^CD44^+^CD25^-^CD122^-^Lin^-^), DN2a (CD117^(hi)^CD44^+^CD25^+^CD122^-^Lin^-^), or ILC1s (CD122^+^Lin^-^) from the neonatal thymus, stimulated with IL-2, IL-7, IL-15 with or without IL-12+IL-18 DP/T cells (CD3e^+^/CD4^+^/CD8b^+^), DN (CD122^-^Lin^-^), ILC1s gated on (CD122^+^NK1.1^+^Lin^-^). **(E)** Gating on KLRG1 and CD27 expression in CD122^+^NK1.1^+^Lin^-^-gated ILC1s following six days of culture in ETP and ILC1-seeded conditions. Gating on KLRG1 and CD27 positive and negative ILC1s were defined based on the expression in all samples analyzed. **(A)** Data is representative of 9 biological replicates from 3 independent experiments. **(D)** Data are shown as Bars with SD from two technical replicates (*n*=2) and representative of three independent experiments (without IL-12+IL-18) and two independent experiments (IL-12+IL-18-induced KLRG1^+^ ILC1s). **Abbreviations:** Lin^-^ = Lineage negative, defined as TCRβ^-^TCRγδ^-^CD3e^-^CD4^-^CD8β^-^. DN = Double negative, ETP = Early Thymic progenitors, E10.5 = Embryonic day 10.5, ETP = Early Thymic Progenitors, P0 = Postnatal day 0 (day of birth), SP = Single positive.

### Neonatal thymic ILC1s shift expression from CX3CR1 to CXCR6 upon IL-12+IL-18 stimulation

To compare the NTOC-derived ILC1s with other group-1 ILCs, we performed scRNAseq on group 1 ILC-enriched cells from the liver, spleen, bone marrow, and thymus from P0.5 neonates and 8-week-old adult mice. Group-1 ILCs were identified based on their *Ncr1*^+^*Cd3e^-^ phenotype*. All cells co-localizing with the group-1 ILCs (ILC precursors, ILC2s and ILC3s) were included in the downstream comparison. ILC1s from NTOC supernatant (**Figure 1M**) were included along with two internal control clusters (CD4 T cells and yδT17 cells) (**Figure 3A**) to avoid proliferation genes confounding the clustering only non-proliferating cells were included from the NTOC supernatant. After integration, UMAP and marker gene expression revealed six major groups of cells as indicated by color (**Figures 3B, 3C, and S3A)**.

**Figure 3:**
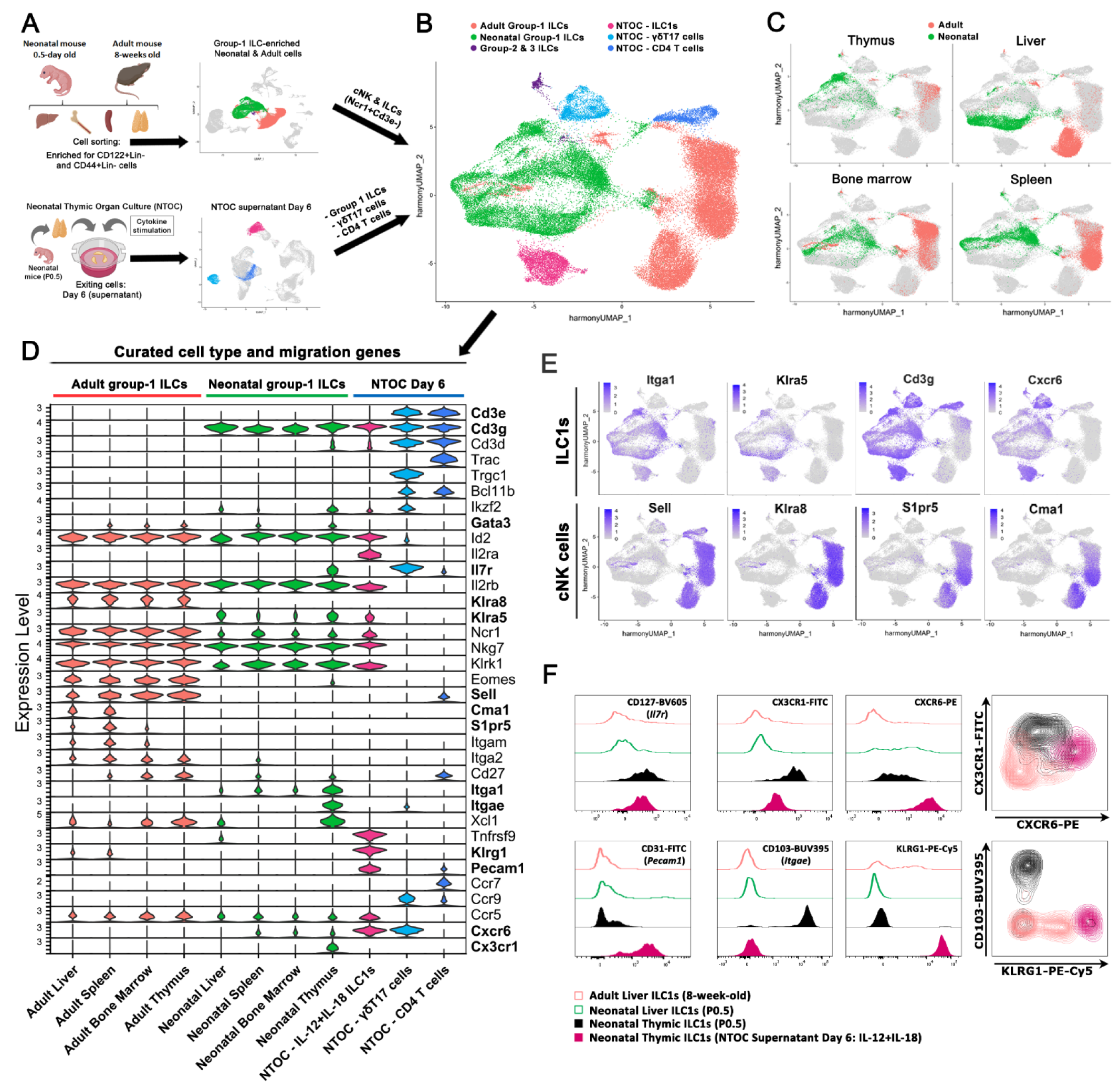
Neonatal thymic ILC1s shift expression from CX3CR1 to CXCR6 upon IL-12+IL-18 stimulation. **(A)** Schematic of scRNA-seq strategy to extract liver, bone marrow, spleen, and thymus from P0.5 neonates and 8-week-old adult mice. Cell enrichment was performed, and CD122^+^Lin^-^ cells and CD44^+^Lin^-^ cells were sorted for unbiased enrichment of group-1 ILCs (ILC1s and cNK cells) and progenitor cells from each of the organs. The resulting eight scRNAseq data sets were combined, and from these, the central grouping of cells primarily containing ILC1s and cNK cells (identified by *Ncr1^+^Cd3e^-^* expression) were selected for downstream analysis. Additionally, three clusters from the scRNAseq data in Figure 2 were selected from the combined four conditions of NTOC supernatant for integration with new single-cell data: **1)** ILC1s (IL-12+IL-18), **2)** γδT17 cells, **3)** and CD4 T cells. The two T cell clusters were chosen as internal controls. **(B)** UMAP of Harmony integrated data explained with the same colors as in **A**, showing the resulting 6 major groupings divided based on sample origin, and in the case of Group-2 & 3 ILCs gene expression. **(C)** UMAP visualization of cells derived from the four tissues green (neonates) and red (adults). **(D)** Violin plots showing gene expression of curated genes with colors based on **B,** showing the group-1 ILCs and the eight different tissues of origin and the three NTOC clusters. **(E)** Feature plots of ILC1 markers (top row) and cNK cell markers (bottom row). **(F)** Flow cytometry comparison of ILC1s (CD122^+^NK1.1^+^CD49a^+^CD62L^-^Lin^-^) from adult the liver (8-weeks-old), neonatal liver and thymus (P0.5), and IL-12+IL-18-stimulated NTOC supernatant (Day 6), displayed as histograms with thymus-derived ILC1s (closed) and liver ILC1s (open) or density plots (showing outliers down to the 2^nd^ percentile). Data are representative of 6 biological replicates (CD127; CX3CR1; CXCR6; CD103; KLRG1) or 3 biological replicates (CD31). **Abbreviations:** Lin^-^: Lineage negative, defined as TCRβ^-^TCRγδ^-^CD3e^-^CD4^-^ CD8β^-^Ter-119^-^Ly-6G^-^F4/80^-^CD19^-^CD11c^-^. NTOC = neonatal thymic organ culture. UMAP = Uniform Manifold Approximation and Projection.

Next, all group-1 ILCs (ILC1s and cNK cells) were divided based on their organ of origin and compared to the three NTOC clusters identified in the cells from the supernatant at day 6 (ILC1s, γδT17 cells, and CD4^+^ T cells). The group-1 ILCs from all organs shared expression of *Id2*, *IL2rb*, *Ncr1*, *Nkg7* and *Klrk1* (**Figure 3D**). Strikingly, there was a clear dichotomy in the expression profile between adult and neonatal-derived group-1 ILCs. The neonatal and NTOC group-1 ILCs expressed the ILC1-specific^19, 36^ markers *Itga1, Klra5*, *Cd3g*, and *Cxcr6*. However, they did not express the cNK cell-specific^36^ markers *Sell*, *Klra8*, *S1pr5*, or *Cma1*, thus reaffirming that neonatal group-1 ILCs primarily consist of ILC1s **(Figures 3D-3E)**. Interestingly, neonatal thymic ILC1s showed high expression of several markers which were not correspondingly expressed in the other tissue-derived group-1 ILCs, such as *Il7r*, *Itgae*, and *Cx3cr1*. In contrast, IL-12+IL-18-activated NTOC ILC1s showed high expression of *Il2ra*, *Klrg1*, *Tnfrsf9*, *Pecam1*, *Cxcr6*, *Ifng*, *Prf1*, *Gzmb*, and *Gzmc* (**Figures 3E** **and S3B)**. Flow cytometry experiments validated these findings and showed that the majority of neonatal thymic ILC1s could be defined as CX3CR1^+^CD103^+^CD122^+^Lin^-^. Conversely, following stimulation with IL-12+IL-18, NTOC-derived ILC1s showed reduced CX3CR1 and no CD103 expression, shifting to the expression of CXCR6 and CD31 (**Figure 3F**). The shift of thymic ILC1s going from *in situ* CX3CR1 expression to CXCR6 expression following IL-12+IL-18 stimulation could indicate a changed homing capacity in response to type 1 cytokine activation.

### Steady-state neonatal thymic ILC1s display a unique and highly immature phenotype

Following the Harmony integration, the UMAP of adult, neonatal and NTOC-derived cells were divided into 20 clusters (**Figures 4A** **and S4A).** Several populations were found to cluster according to cell state rather than cell type or tissue imprinting; Cluster 1 contains dying cells **(Figures S4B and S4C),** cluster 2 contains a high expression of interferon-stimulated genes (ISG) **(Figure S4A)**, and clusters 9 and 11 comprise proliferating cells **(Figure S4A).** To further focus the downstream analysis on the cell type and tissue-specific imprinting, we selected the remaining eleven group-1 ILC clusters and one cluster of ILC precursors **(Figures 4B, 4C, S4A, and S4D).** Based on the gene expression in the different clusters, we divide ILC1s into Helper-like and Cytotoxic ILC1s (**Figure 4D**). As shown in the perinatal liver^19, 36^, the neonatal liver group-1 ILCs (Cluster 12) mainly comprised cytotoxic ILC1s. A small population were identified as adult ILC1s (Cluster 17) with higher gene expression similarity to neonatal ILC1s than adult cNK cells. Interestingly, the neonatal thymic ILC1s (Cluster 18) displayed an immature helper-like phenotype and expressed higher levels of associated markers (*Tox, Tcf7, Cd27,* and *Il7r*), than other neonatal ILC1 clusters (**Figures 4E** **and S4D)**. Accordingly, analysis of group-1 ILC transcription factors revealed that the thymic ILC1s and cNK cells expressed the highest levels of stem-like markers^36, 40^ such as *Tcf7*, *Batf3, and Tox* among group-1 ILC clusters, along with high expression of Hobit (*Zfp683*), *Gata3*, and *Ikzf2* in the thymic ILC1s compared to the other ILC1 clusters **(Figure S4E)**. The vast majority of the adult group-1 ILCs were identified as cNK cells. Based on previous single-cell studies^19^, the genes (*Tcf7*, *Cd27* and *Emb*) decreased while the genes (*Itgam*, *Klrg1*, *Prf1*, *Gzmb, Zeb2*, *Cma1*, and *S1pr5,*), increased according to cNK maturation level. They were therefore used to divide cNK cells into early, mid, and late developmental stages (**Figures 4E** **and S4D).** The Ly49 family genes (*Klra1* - *Klra8*) allowed the separation of the adult group-1 ILCs and neonatal ILC1s **(Figure S4F)**, confirming previous results showing *Klra5* as a specific embryonic-wave ILC1 marker, indicating neonatal thymic ILC1s deriving from fetal liver precursors^19, 20^. In conclusion, the steady-state thymus primarily contains ILC1s with a unique embryonic-wave phenotype and displays higher levels of helper-like ILC1 genes than other neonatal ILC1 clusters.

**Figure 4:**
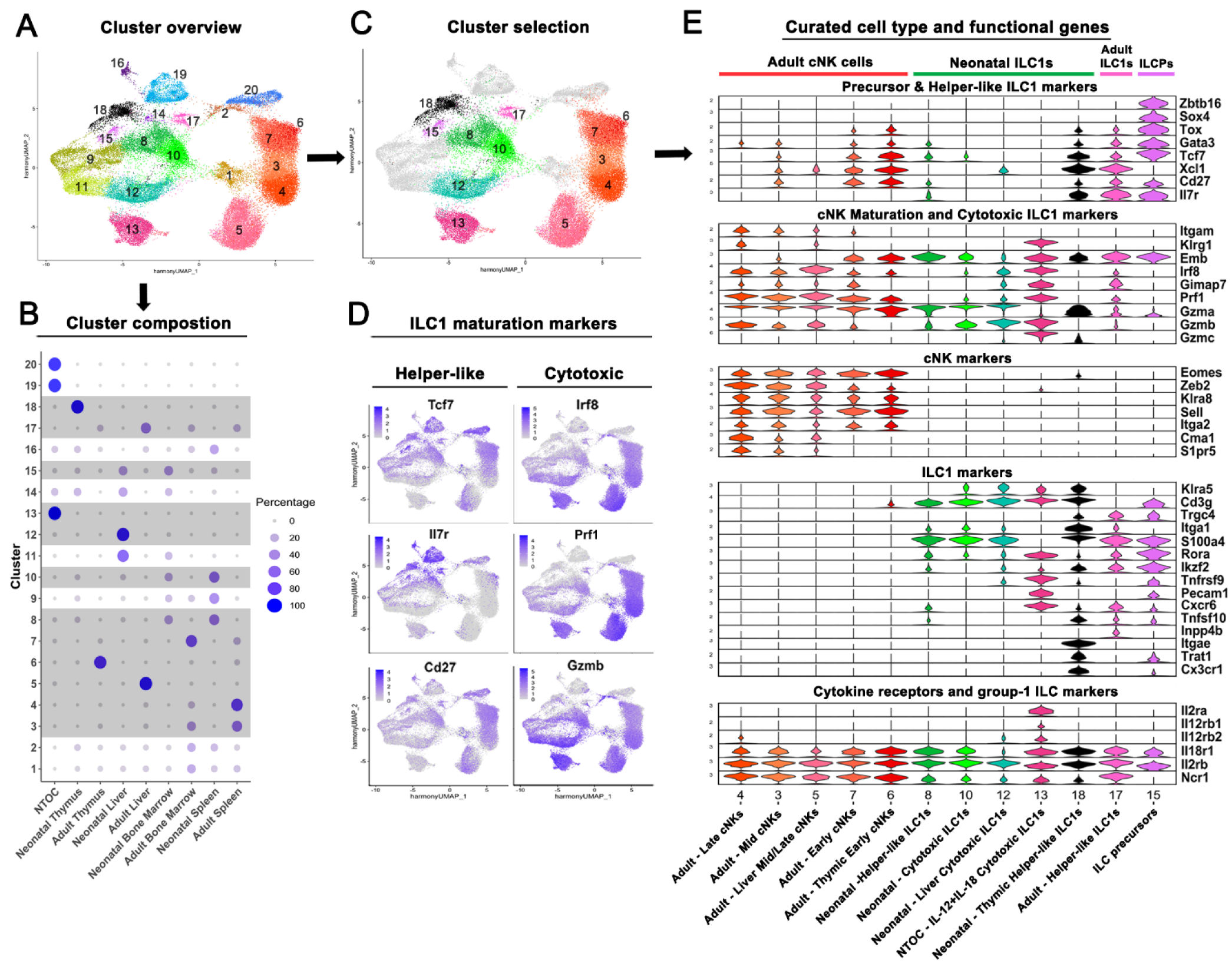
Neonatal thymic ILC1s have a unique phenotype *ex vivo*. **(A)** UMAP overview of all 20 clusters in Harmony-integrated scRNAseq comparison of group-1 ILCs in 8-weeks-old adult and P0.5 neonatal mice across tissues (Thymus, Liver, Spleen, Bone marrow) and NTOC clusters. **(B)** Cluster composition overview, showing the tissue distribution of each of the 20 clusters, with the selected clusters from **C** shown with gray background. **(C)** UMAP overview of selected clusters in group-1 ILC comparison. **(D)** Feature plots showing gene signatures associated with Helper-like ILC1s (Left column), and Cytotoxic-like ILC1s genes (Right column). **(E)** Violin plot showing gene expression of curated group-1 ILC genes in selected clusters. Cluster order is based on similarity in visualized gene expression between clusters. **Abbreviations:** ILCPs = Innate lymphoid cell precursors. NTOC = neonatal thymic organ culture. UMAP = Uniform Manifold Approximation and Projection.

### *In situ* imaging of ILC1s reveals expansion in neonatal thymus following MCMV infection

The *Ncr1*^iCre^ and Rosa26-tdTomato mouse lines were crossed to create “*Ncr1*-tdTomato” mice, used for group-1 ILCs fate-mapping^41^. Thus, enabling the visualization of the expanding ILC1s in the neonatal thymus. We performed confocal 3D imaging of the entire thymic lobe. Following MCMV infection on P2, an expansion of *Ncr1*-tdTomato^+^ cells was observed in the cortex and capsular region of the thymus five days after (P7). This expansion did not occur in sham control thymic lobes. Furthermore, most *Ncr1*^+^ cells did not show overlap with CD3e-expressing cells **(Figure 5, rows 1-2)**. Similarly, NTOCs using *Ncr1*-tdTomato thymic lobes revealed enhanced production of Ncr1^+^ cells on Day 3 following exposure to IL-12+IL-18 stimulation compared to the vehicle condition **(Figure 5, rows 3-4)**. These data substantiate the previous results by both showing ILC1 expansion during type 1 immunity.

**Figure 5:**
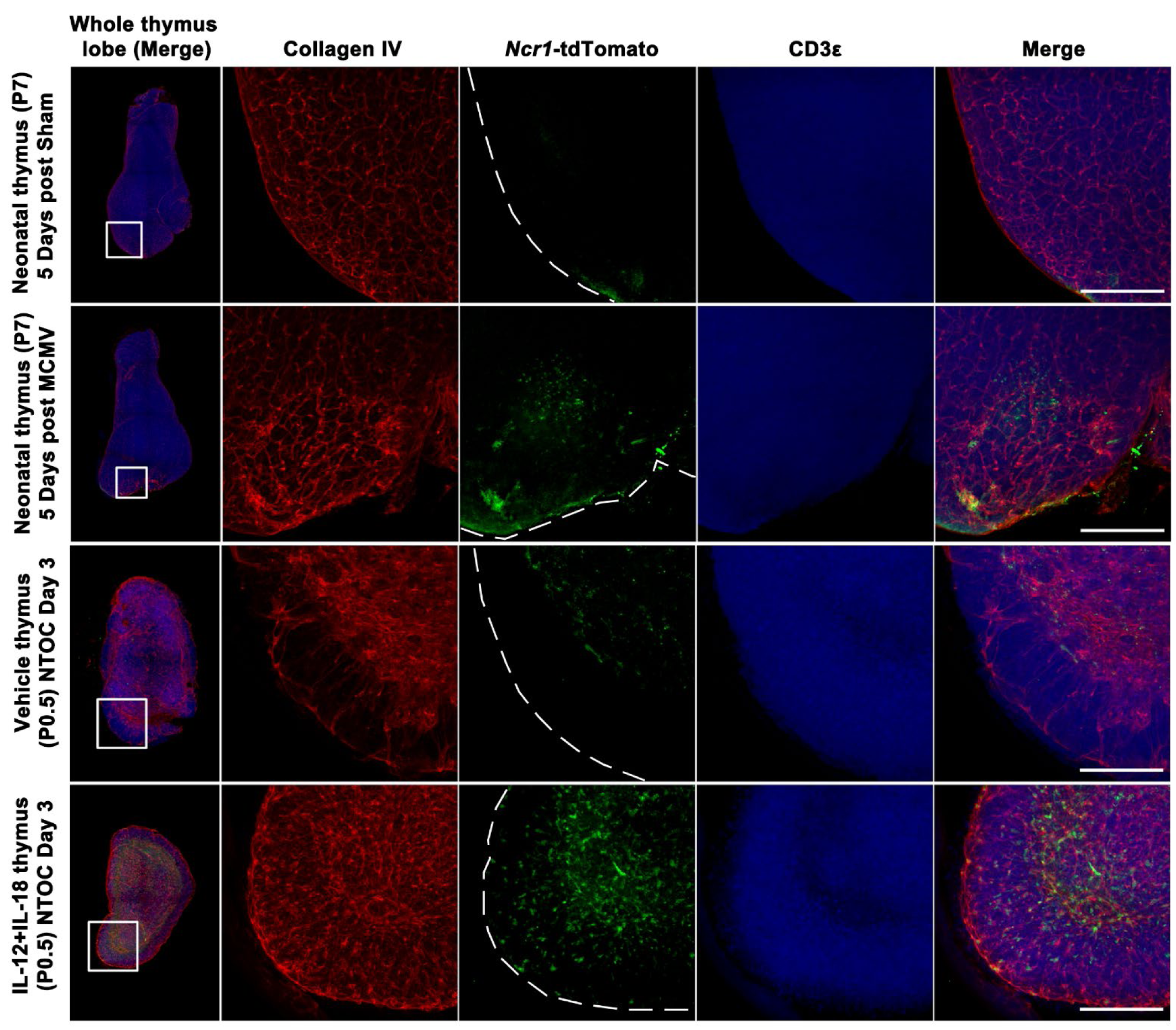
In situ imaging shows altered localization of neonatal thymus ILC1s during MCMV infection. **(Column 1)** Confocal 3D imaging of neonatal thymic lobes expressing Ncr1-tdTomato. A whole thymic lobe imaged showing the zoomed-in section used in the other four columns. **(Column 2-4)** Single colors and **(Columns 1 and 5)** merged overlay of all colors are shown as maximum projection intensity: Red (Collagen IV), Green (Ncr1-tdTomato), and Blue (CD3). **(Rows 1 and 2)** Neonatal thymus from Sham or MCMV infected mice was extracted 5 days after infection (injected intraperitoneally with 3000pfu on P2). **(Rows 3 and 4)** Thymic lobes from Day 3 vehicle or IL-12+IL-18 stimulated NTOC. The bars in the right corner are all 200 μm. Images are representative of 3-5 biological replicates (Sham: *n*= 3; MCMV: *n*=5; Vehicle NTOC: *n*=3; IL-12+IL-18 NTOC: *n*=4) from two independent experiments. **Abbreviations:** NTOC = neonatal thymic organ culture. MCMV = Murine cytomegalovirus.

### KLRG1^+^ thymic ILC1s home to the liver and the peritoneal cavity

The NTOC cultures revealed ILC1 exit and expansion from thymic lobes as well as high CXCR6 expression, which may indicate a capacity for liver homing^36, 42^. Thus, we wanted to document whether exit and expansion would be associated with homing of thymus-derived ILC1s to peripheral compartments *in vivo*. Thus, we established a thymic kidney capsule implantation model^43^, where P0.5 neonatal thymic lobes from *Ncr1*-tdTomato mice were grafted under the kidney capsule of *Rag2*^-/-^*Il2rg*^-/-^ mice, which lack T cells, B cells and ILCs. Thus all T cells and tdTomato^+^ ILC1s observed in these transplanted mice are derived from the neonatal thymus graft. IL-2 is critical for the IL-12+IL-18-synergy^44, 45^, and due to their lack of T cells, *Rag2*^-/-^*Il2rg*^-/-^ mice have severely impaired IL-2 production^46^. Therefore, IL-2 was either administered alone, as a control, or in combination with IL-12+IL-18. Following three days of vascularization of the grafted thymus, the mice were injected with cytokines IP every 24 hours for a further three consecutive days, and mice were analyzed on Day 6 after grafting (**Figures 6A** **and 6B)**. We examined the thymus graft, spleen, liver, and peritoneal cavity by flow cytometry for the presence of thymic graft-derived T-cells and tdTomato^+^ ILC1s. We confirmed the successful grafting by repopulating CD4^+^ and CD8^+^ T cells in the spleen **(Figure S6A)**. We further observed a substantial influx of neonatal tdTomato^+^ ILC1s into the liver, peritoneal cavity, and spleen after administration of IL-2. Interestingly, only homing of ILC1s in the liver and peritoneum was enhanced by the administration of IL-12+IL-18 compared to IL-2 injected mice (**Figures 6C** **and S6B)**. This reveals that thymic-derived ILC1s develop and are exported in the absence of type-1-induced inflammation. In both conditions, low numbers of ILC1s remained in the thymic graft, indicative of the thymic ILC1 exit **(Figure S6C)**. Moreover, we observed that the homing of thymus-graft-derived CD4 T cells was not significantly changed by the different conditions **(Figure S6D)**. Thus, to evaluate the KLRG1^+^ ILC1s homing efficiency in the six days following the grafting, we determined the ratios of CD27^+^KLRG1^-^ and KLRG1^+^ ILC1s to the amount of CD4 T cells in each organ. Both CD27^+^KLRG1^-^ and KLRG1^+^ ILC1s showed a much higher homing to the peritoneal cavity and liver than the spleen. However, CD27^+^KLRG1^-^ ILC1s did not show any difference in homing capacity between conditions, whereas in the IL-2+IL-12+IL-18-condition KLRG1^+^ ILC1s and CD4 T cells entered the peritoneal cavity and the liver approximately in a 1:1 ratio and a 1:2 ratio, respectively **(Figures 6D, 6E, S6E, and S6F)**. The KLRG1^+^ ILC1s thereby displays a high preferential ILC1-homing to these two compartments during type 1 immunity. This indicates that the IL-12+IL-18-induced KLRG1^+^ ILC1s differentiation results in an increased capacity for tissue homing from the thymus. Finally, to visualize the thymus-derived Ncr1-tdTomato exit and homing to the periphery, we performed confocal imaging on the liver of these mice. Similar to flow cytometry results, we found a clear homing and tissue localization of *Ncr1*-tdTomato ILC1s inside the liver, which was exacerbated in mice receiving IL-12+IL-18 (**Figure 6I**).

**Figure 6:**
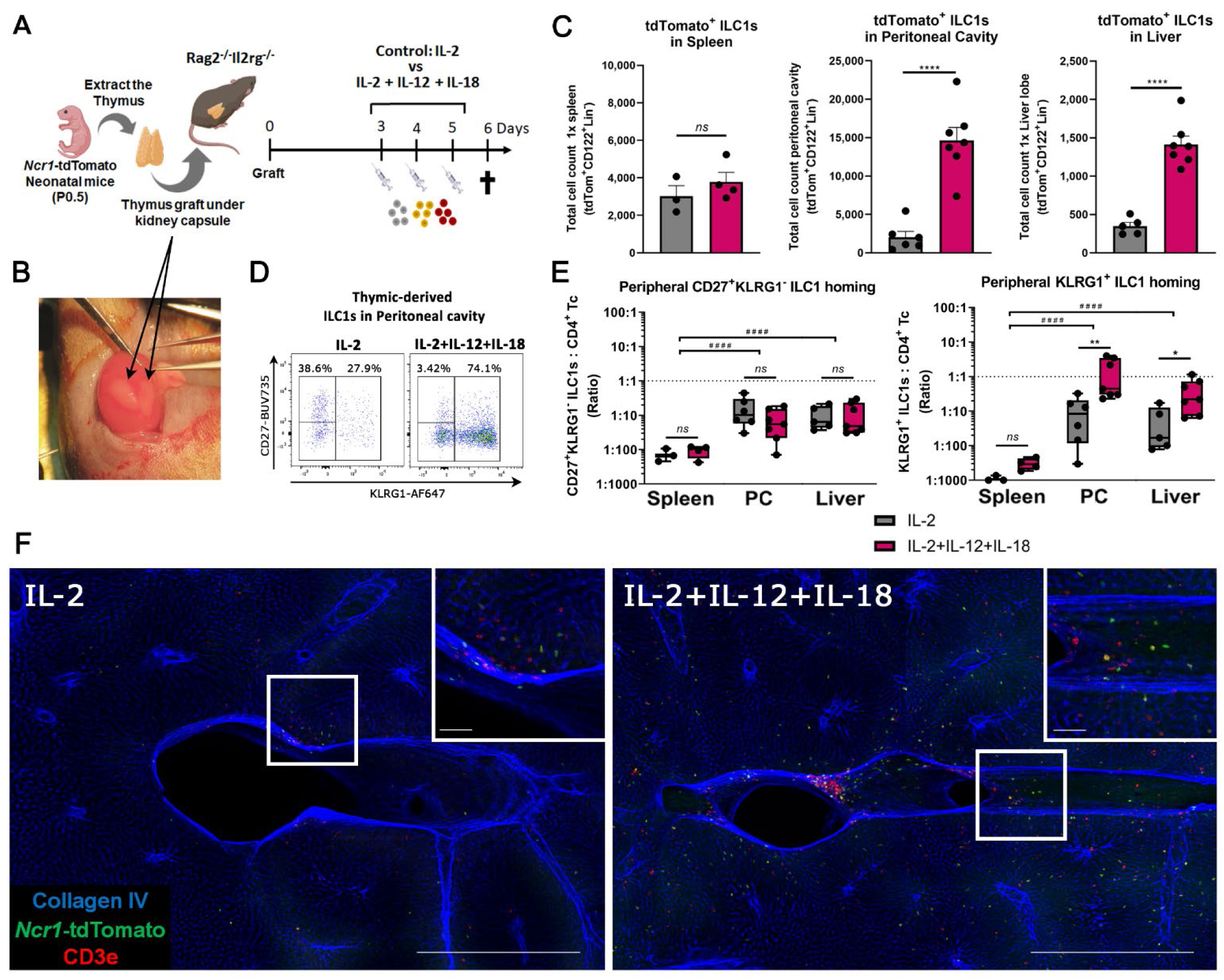
IL-12+IL-18 activated thymic ILC1s home to the liver and peritoneal cavity. **(A)** Schematic of neonatal thymus graft experiment, where *Ncr1*-tdTomato thymus was grafted under the kidney capsule of in *Rag2*^-/-^*Il2rg*^-/-^ deficient mice. From Day 3 after grafting, mice were injected three times with either IL-2 (control) or IL-2+IL-12+IL-18 with 24h in between each injection. Mice were terminated six days after grafting. **(B)** Picture of the kidney with two recently grafted neonatal thymic lobes. **(C)** Bar plots of thymus-graft-derived tdTomato^+^ ILC1s in the spleen, peritoneal cavity, and liver. **(D)** Flow plots showing the gating strategy of KLRG1 and CD27 in thymus graft-derived tdTomato-expressing ILC1s in two conditions, shown from the peritoneal cavity. **(E)** Box plots, showing the count *ratio* on a Log10-scale between thymus-graft-derived ILC1s and CD4 T cells in the spleen, peritoneal cavity (PC), and liver, **Left**: CD27^+^KLRG1^-^ILC1s : CD4 T cells (Ratio), and **Right**: KLRG1^+^ ILC1s : CD4 T cells (Ratio). **(F)** Confocal 3D imaging of 500μm thick liver samples from Ncr1-tdtomato thymus-grafted mice six days after grafting, representative pictures for each condition are shown as maximum projection intensity, Blue (Collagen IV), Green (*Ncr1*-tdTomato), Red (CD3), the bar in the bottom right corner is 500 μm, and bar in left corner of zoomed section is 50 μm, representative of three-four biological replicates (IL-2: *n*=4; IL-2+IL-12+IL-18: *n*=3). **(C)** Error bars are shown as SEM and **(E)** Box and whiskers plots with Min and Max, displaying median and 25^th^ to 75^th^ percentiles. **(C and E)** each dot represents a separate thymus-grafted mouse (spleen: *n*=3-4; peritoneal cavity: *n*=6-7, Liver: *n*=5-7), and data are combined from two independent experiments. Statistics; **(C)** F test showed equal variances for treatment found no difference in variance between treatments. **(E)** F test showed equal variances on the log2-transformed scale for the conditions. Thus differences were evaluated by Two-anova (Using Sidak’s Multiple comparisons test) on log2-transformed data, which showed significant differences between the two treatments for PC ((effect size (4.133); t-value (4.095); df (26); Adjusted *p*-value (0.0011)), and for Liver ((effect size (3.167); t-value (2.990); df (26); Adjusted *p*-value (0.0180)). There was no significant difference between the treatments for the spleen ((effect size (2.147); t-value (1.549); df (26); Adjusted *p*-value (0.3492)). **(C)** Unpaired T test (two-tailed) and **(E)** Two-way anova between treatments: * p < 0.05, ** p < 0.01, **** p < 0.0001, *ns* = Not significant. **(E)** Two-way anova between tissues: ^#^ ^#^ ^#^ ^#^ p < 0.0001. **Abbreviations:** Lin-: Lineage negative, defined as TCRβ^-^TCRγδ^-^CD3e^-^CD4^-^CD8β^-^Ter-119^-^Ly-6G^-^. PC **=** peritoneal cavity, Tc = T cell.

## Discussion

Group-1 ILCs can develop from early thymic progenitors (ETPs) in mice and humans^24–27^, though the relationship between thymic group-1 ILCs and peripheral group-1 ILCs remains unresolved^23^. To illuminate unexplored areas of group-1 ILC ontogeny, Sparano et al. (2022) used fate-mapping and scRNAseq by describing different embryonic and neonatal waves of ILC1s and cNK cells in the liver and bone marrow, showing that the vast majority of group-1 ILCs are ILC1s rather than cNK cells at birth^19^. Our scRNAseq data shows that ILC1s from the neonatal thymus display unique markers compared to ILC1s from bone marrow, spleen, or liver, identifying them as a distinct ILC1 subset. Interestingly, most neonatal thymic ILC1s exhibit embryonic-wave characteristics (*Klra5*-expression) and an immature helper-like ILC1 phenotype (*Tcf7^(hi)^Il7r^(hi)^Tox^+^Cd27^+^*)^19, 20, 36^. As many aspects of ILC1 ontogeny remain unknown, this finding opens the possibility for some subsets of embryonic-wave-seeded peripheral ILC1s to originate from the perinatal thymus.

In parallel to ILC1s, ILC2s can also develop from progenitors in the thymus^47, 48^. A study on the intrathymic transcriptional checkpoints revealed that Rora expression in embryonic thymic progenitors repressed T cell fate and promoted thymic ILC2 commitment^47^. However, although *Rora* expression was highly increased in the thymus-exiting ILC1 population, it remains unknown whether thymic ILC1s have an analogous *Rora* dependency as thymic ILC2. Similar to previous findings^24, 26^, we show that perinatal ETPs have a substantial group-1 ILC potential, even in T cell-skewing OP9-DL1 conditions. Furthermore, we find that the developing group-1 ILCs are ILC1s rather than cNK cells, as previously believed. These ILC1s lack the differentiation maker KLRG1 following six days of culture, despite the presence of IL-12+IL-18 in the culture. Only after seeding immature neonatal thymic-derived ILC1s and stimulating with IL-12+IL-18 do the ILC1s acquire the terminal differentiation marker KLRG1^38^, similar to the NTOC-exiting ILC1s. These data indicate that the expanding cytotoxic ILC1s observed in the NTOC supernatant mainly derive from the immature ILC1s already present in the neonatal thymus rather than inducing their *de novo* development from the ETPs. Although group-1 ILCs can develop from human thymic progenitors following *in vitro* culture, direct *in situ* group-1 ILC development in the human thymus has yet to be confirmed^25, 27^. Whether infection or inflammatory triggers are required to perturb steady-state thymic development to allow *in situ* group-1 ILC development remains to be determined.

Upon activation with IL-12+IL-18, the ILC1s downregulated CX3CR1 and upregulated CXCR6 and CD31. CXCR6 expression on ILC1s is associated with liver homing^36, 42^, and CD31 has been associated with trans-endothelial migration in multiple immune subsets^49^, which may be important for entering peripheral tissues. Thus, the shift from CX3CR1 to CXCR6/CD31 expression reflects an apparent change in the homing capacity of the thymic-derived ILC1s. Accordingly, we used subcapsular thymic kidney grafts to track ILC1s from the P0.5 neonatal thymus. Despite ILC1s being a rare population in the thymus, a substantial ILC1 population was observed in all examined tissues six days after grafting, indicative of more peripheral ILC1 homing from the thymus graft than can be explained by the small number of thymic ILC1s present at birth. In addition, we compared the number of ILC1s and CD4 T cells entering the tissues and surprisingly found that the presence of thymic graft-derived ILC1s and CD4+ T cells in the liver and peritoneal cavity were relatively similar in both tissues. However, unlike the liver and peritoneal cavity, the amount of thymic-derived ILC1s was 75-120 times lower in the spleen compared to CD4 T cells. These results and the proportional rarity of thymic ILC1s indicate a preferentially homing of thymic ILC1s towards the liver and the peritoneal cavity. TCRa-KO thymus grafts were previously used to show group-1 ILCs homing to the spleen^22^. However, to our knowledge, peripheral homing of thymic group-1 ILCs has not been shown in wild-type thymus grafts. Thymic-derived ILC1 homing to the liver and peritoneum increased substantially in response to IL-12+IL-18 compared to IL-2-injected control mice and consisted primarily of differentiated KLRG1^+^ ILC1s.

Although the underlying causality for infection-induced thymus atrophy is poorly understood^11^, infection with murine roseolovirus has been shown to disrupt central tolerance in the thymus^50^. Here we show that acute thymus atrophy is not only a process where developing T cells passively die but that type-1 inflammation actively mediates *in situ* thymic ILC1 expansion. Our results contribute to existing evidence that the thymus is not only a T cell-producing organ but can also produce peripheral homing ILC1s, which is augmented during acute type-1 inflammation. Although we do not definitively show that embryonic ILC1s derive from the thymus, this work is the first to show that peripheral ILC1s can derive from the thymus and that some subsets of embryonic-wave ILC1s are likely to have a thymic origin.

In conclusion, we show that MCMV infection and IL-12+IL-18-induced sterile inflammation result in thymus atrophy in neonatal mice. *Ex vivo* culturing of neonatal thymic lobes with IL-12+IL-18 results in a substantial expansion and thymic exit of KLRG1^+^ ILC1s. We confirm that these thymus exiting KLRG1^+^ ILC1s have an embryonic wave phenotype and differentiate from immature CX3CR1-expressing thymic ILC1s. NTOC-derived ILC1s show enhanced CXCR6 expression following IL-12+IL-18 stimulation. Furthermore, 3D confocal imaging shows ILC1 expansion in the thymus of MCMV-infected neonates. Finally, using fate-mapping neonatal thymus grafts, we found that ILC1s could exit the neonatal thymus to the periphery and that type-1 inflammation enhanced the preferential homing of thymic-derived KLRG1^+^ ILC1s to the liver and peritoneal cavity. The results reveal a novel thymic function by contributing to type 1 immune responses through the rapid production of peripheral homing ILC1s.

## Author contributions

P.T. and P.V. supervised the project. P.T., G.L., and P.V. designed experiments. P.T., M.R., W.S., T.D., and J.V., performed experiments. J.H. optimized the cytotoxicity assay. J.V, optimized MCMV batch creation under the supervision of S.J.. T.T., T.B.B. N.T., S.J., and G.L., contributed to discussions and critical data interpretation. P.T., and M.R. created the illustrations. A.G. was instrumental for bioimaging support on 3D imaging and analyzed the imaging data. B.V., R.R., and T.B.B. performed scRNA-seq analyses. P.T., M.R., and B.V. analyzed data. P.T. wrote the manuscript. P.V., G.L., and T.B.B., provided considerable input to the manuscript. All of the authors have read or provided comments on the manuscript.

## Acknowledgements

We thank the VIB Single Cell Core, VIB Nucleomics core, VIB Flow Core Ghent and VIB Bioimaging core for support and access to the instrument park (vib.be/core-facilities). We want to thank Søren Skov (UCPH-SUND), Jonathan Maelfait (VIB-UGent), Kodi Ravichandran (VIB-UGent), Andrew Brown (VIB-UGent), and Bart Lambrecht (VIB-UGent) for their critical input to the project. We are grateful to Eric Vivier for providing us with the Ncr1-iCre mouse line and Bart Lambrecht for giving us the *Rag2*-/- OTI and the *Rag2*-/-*Il2rg*-/- mouse lines. We thank Bastiaan Maes (VIB-UGent) and Jonathan Maelfait (VIB-UGent) for their feedback on the manuscript. Finally, we thank the VIB-UGent animal house staff. Biorender was used to create some figures.

## Online Methods

### Mice and Ethical considerations

C57BL/6N mice were used in all experiments except for the transgene mice and their wild-type controls, which were on a C57BL/6J background. Transgene mice: Ncr1^iCre^ mice (Ncr1^tm1.1(icre)Viv^) were generously provided by E. Vivier^41^ and crossed with ROSA26–flox-stop-flox-tdTomato mice (B6.Cg-Gt(ROSA)26Sortm14(CAG-tdTomato)Hze/J) to create Ncr1-tdTomato mice. Rag2^-/-^OTI (B6.129S6-*Rag2^tm1Fwa^* Tg(TcraTcrb)1100Mjb), and Rag2^-/-^Il2rγ^-/-^ mice (C;129S4-*Rag2^tm1.1Flv^ Il2rg^tm1.1Flv^*/J). Purchased time-mated pregnant females were bought from Janvier. All transgene mice were bred and kept in-house at the VIB Center for Inflammation Research under specific pathogen-free (SPF) under specific pathogen-free (SPF) conditions. All mice were kept in individually ventilated GM500 cages (Techniplast) with a 14/10-hour light/dark cycle in a temperature-controlled (21°C). Mice were provided water and food ad libitum. All mouse experiments were conducted according to institutional, national, and European animal regulations. Ghent University’s Ethics Committee approved all experimental animal procedures. Experiments were conducted in agreement with the European Parliament’s Directive 2010/63/EU and the 22-09-2010 Council on the protection of animals used for scientific purposes.

### Genotyping

Genotyping of the different transgene mouse lines was performed by PCR. *Ncr1-tdTomato* mice: tdTomato^fl/fl^ mice were genotyped using the PCR primers: wild-type forward primer CAGCTTCTTGTAATCGGGGA, wild-type reverse primer GCGAGGAGGTCATCAAAGAG, Transgene forward primer TGGCTTCTGAGGACCGCCCTGGGC; Transgene reverse primer CACCTGTTCAATTCCCCTGCA. Yielding a 160bp wild-type DNA band is at 160bp and a transgene DNA band of 245bp. Ncr1^iCre^ mice were genotyped using the PCR primers: Forward-1 primer ACCGCCTGATCTATGGTGCC, Forward-2 primer GGGTGGGTGTAGCCTCTATC, Reverse primer CACTCCTACCCCTTCATTTCTGA. Yielding a 350bp wild-type DNA band and a transgene DNA band of 280-300bp. *Rag2^-/-^OT-I mice* (B6.129S6-*Rag2^tm1Fwa^* Tg(TcraTcrb)1100Mjb) were genotyped using the PCR primers. Rag2^-/-^: Forward-1 primer CAGGGTCGCTCGGTGTTC, Forward-2 primer CTTGCCAGGAGGAATCTCTG, Reverse primer GTTTCCCATGTTGCTTCCA, yielding a 200bp wild-type DNA fragment and a 388bp knockout DNA fragment. OT-I: OT-I TCR Forward CAGCAGCAGGTGAGACAAAGT, OT-I TCR Reverse GGCTTTATAATTAGCTTGGTCC, OT Int Ctrl Forward CAAATGTTGCTTGTCTGGTG, OT Int Ctrl Reverse GTCAGTCGAGTGCACAGTTT. Yielding a 200bp wild-type DNA band and a 300bp transgene DNA band. Rag2^-/-^Il2rγ^-/-^ mice were genotyped using the same Rag2^-/-^ primers described above and Il2rγ^-/-^: Common Forward GTGGGTAGCCAGCTCTTCAG, wild-type reverse CCTGGAGCTGGACAACAAAT, and knockout reverse GCCAGAGGCCACTTGTGTAG. Yielding a 269 bp wild-type DNA fragment and a 349 bp knockout DNA fragment.

### Cytokines

The following cytokines were used: Recombinant murine IL-2 (Protein core VIB, Ghent, Belgum). Recombinant murine IL-7 (Cat# 217-17, Peprotech, Rocky Hill, NJ). Recombinant murine IL-12 (Cat# 210-12, Peprotech, Rocky Hill, NJ). Recombinant murine IL-15 (Cat#, 210-15, Peprotech). Recombinant murine IL-18 (Cat# P70380, R&D Systems, Minneapolis, MN). All lyophilized cytokines were reconstituted in PBS with 0.1% BSA before storing at −80°C. The concentration of BSA in all vehicle control conditions corresponded to the highest concentration within the specific experiment.

### Neonatal thymic organ cultures (NTOC)

Neonates were euthanized by decapitation at P0.5 and thymic lobes were extracted, and cleaned before setup. Three neonatal thymus lobes were cultured per well unless otherwise stated. Only intact neonatal thymic lobes were cultured on top of a 0.8 μm IsoporeTM Membrane Filter (Cat# ATTP01300, Merck Millipore), which was suspended on top of 2 mL NTOC medium in 12-well plates and cultured at 37°C with 5% CO_2_. Cytokines were used at the following concentrations: IL-12 [4 ng/mL], and IL-18 [40 ng/mL]. The NTOC medium was composed of DMEM (Cat# 31330-038, GIBCO) supplemented with 20 % FCS (Lot# 71133 and 90439, Brand: TICO EUROPE, Greiner and Int Medical), 20μM L-glutamine, 50 μM β-mercapto-ethanol, 0.8 % penicillin-streptomycin (#Cat P4333, Sigma-Aldrich), 0.4 mM Na pyruvate (#Cat S-8636, Sigma-Aldrich), 1x non-essential amino acids (#Cat M7145, Sigma-Aldrich). Different types of FCS were tested for optimal cellular output of the NTOC to ensure it supported ILC1 development.

### OP9-DL1 co-cultures

1,000 OP9-DL1 engineered bone marrow stromal cells were plated in 96 well plates 24 hours prior to co-culture. We sorted the population of interest: Early thymic progenitors (ETPs) [CD44^+^CD25^-^c-Kit^high^Lin^-^], DN2a [CD44^+^CD25^+^c-Kit^high^Lin^-^] and immature ILCs [CD44^+^CD122^+^Lin^-^] in a Symphony A5 FACS sorter (BD Biosciences). Co-cultures were initiated by seeding 1,000 cell-sorted precursors/immature ILC1s cells were seeded and treated with cytokines. Cytokines were used at the following concentrations: IL-2 [20 ng/mL]; IL-7 [20 ng/mL]; IL-15 [20 ng/mL], IL-12 [4 ng/mL], IL-18 [40 ng/mL] in a final volume of 200µL. The OP9-DL1 cell line and co-cultures were maintained at 37°C with 5% CO2, in αMEM (Cat#: 22571-020, GIBCO Gibco) medium containing 20% FCS (TICO), 20μM L-glutamine, 50 μM 2-mercaptoethanol, 0.8 % penicillin-streptomycin (Cat#: P4333, Sigma-Aldrich, P4333), 0.,4 mM Na-pyruvate (Cat#: S-8636, Sigma-Aldrich, S-8636), 1x non-essential amino acids (Cat#: M7145, Sigma-Aldrich, M7145). At Day 7 the outcome was analyzed by flow cytometry in a Symphony A5 (BD Biosciences) device.

### YAC-1 killing assay

YAC-1 target cells were stained with eFluor 670 dye (Cat# 65-0840-85, ThermoFisher) and 5000 stained YAC-1 cells were seeded in each well of a 96-well U-bottomed plate. Effector cells were sorted from IL-12+IL-18 NTOC supernatant; Group-1 ILCs (CD122+Lin-), and from Vehicle NTOC; non-cytotoxic cellular controls (CD8β^-^CD122^-^). An increasing number of effector cells were added to the wells with the freshly seeded YAC-1 cells to create different Effector:Target ratios with a constant amount of target cells. Cells were centrifuged (300g, 2min, RT) immediately after both effector and target cells were added and incubated at 37°C with 5% CO_2_. Dose-dependent killing was determined by flow cytometry following 4 hours of co-culture, by DAPI staining (0.1 µg/mL) and counting the percentage of dying DAPI^+^ cells. YAC-1 and co-cultured cells were maintained in RPMI medium (#Cat R1780-500ML, Sigma-Aldrich) containing 10% FCS (Grenier), 50 μM 2-mercaptoethanol, 0.8 % penicillin-streptomycin (#Cat P4333, Sigma-Aldrich) and split at least two times before seeding.

### Neonatal thymus grafts

The neonatal thymic grafts were performed as previously described^43^. Adult Rag2^-/-^Il2rγ^-/-^ male mice were provided analgesia in drinking water the day before the grafting procedure. Mice were anaesthetized in an isoflurane box and placed in an isoflurane gas mask on a 37° heat plate. The mice were shaven on the left dorsal side at the height of the kidney and disinfected with 70% ethanol, followed by iso-betadine and another round of 70% ethanol. A 1.5 cm longitudinal incision in skin and muscle was made at the height of the right kidney, and the kidney was exposed. A small horizontal incision was made in the kidney capsule on the caudal kidney pole with a 18G needle. A subcapsular space was created underneath the kidney capsule by gently moving a blunt forceps underneath the capsule in the presence of sterile PBS. Subsequently, two thymus lobes from neonatal Ncr1-tdTomato mice were inserted in the subcapsular space and gently pushed to the cranial pole of the kidney. The grafted kidney was placed back in its original location, the muscles were sutured, and the skin was closed with clips and disinfected with iso-betadine. The mice recovered, and the grafted thymus was allowed to vascularize to the ample kidney vasculature for three days. The thymus-grafted mice then received three intraperitoneal injections of cytokines or vehicle with a 24-hour interval. The mice were euthanized six days after grafting.

### Cytokine injections

Adult and neonatal mice were randomly assigned to their group. All mice received three intraperitoneal injections of cytokines or vehicle with a 24-hour interval. Neonates were injected intraperitonally with cytokines diluted in DMEM in a total volume of 20µl using a 29G needle. The needle opening was facing towards the animal upon injection to avoid abdominal fluid exiting during the procedure. In neonates, the cytokine concentrations per injection were: 10 ng [IL-12]; 100 ng [IL-18]. The cytokine injection in adult C57BL/6N mice was modified from a systemic inflammatory IL-12+IL-18-model in BALB/c mice^51^. In the adult thymus-grafted Rag2^-/-^Il2rγ^-/-^ mice, the model was adapted to an IL-2+IL-12+IL-18 model, considering the lacking IL-2 production in Rag2^-/-^Il2rγ^-/-^ mice^46^. For all adult mice, the cytokines were diluted in, PBS and each mouse was injected intraperitoneally with a total volume of 0.2mL. In adult mice, the cytokine concentrations per injection were: 100.000 IU [IL-2]; 150 ng [IL-12]; 750 ng [IL-18].

### Neonatal Murine Cytomegalovirus (MCMV) infection

C57BL/6N and Ncr1-tdTomato mice neonates were injected once intraperitoneally with 10 µL of MCMV diluted in DMEM (Cat# 31330-038, GIBCO) on postnatal day 3 (P3) using 0.5 mL 29G BD Microfine^TM^ (Cat# 324824, BD Biosciences). The needle opening was facing towards the animal upon injection to avoid abdominal fluid exiting during the procedure. The MCMV batch was obtained by infecting MCMV susceptible BALB/c mice and harvesting salivary glands as described^52^. Mice were either Sham injected (DMEM) or infected with a sublethal dose 300 pfu/μL (3000pfu in total). To avoid cross-infection, the infected neonates and sham neonates were caged separately with their biological mothers. All MCMV-infected mice were kept in a BL2 animal facility in the VIB-IRC, and all handling of the virus and tissues were performed in a BL2 facility.

### Electron microscopy

Thymic CD4+ T cells (TCRb+CD4+CD8b-), and ILC1s (CD122+CD4-CD8b-TCRb-TCRgd-CD19-Ly-6G-MHCII-CD11c-) were sorted from NTOC supernatant Day 6 treated with vehicle and IL-12+IL-18 respectively. Sorted cells were fixed in 4% PFA and 2.5% glutaraldehyde in 0.1 M NaCacodylate buffer, pH 7.2 and centrifuged at 1500 rpm. Low melting point-agarose was used to keep the cells concentrated for further processing. Cells were fixed for 4h at RT followed by fixation O/N at 4°C after replacement with fresh fixative. After washing in buffer, they were gaardpost-fixed in 1% OsO4 with 1.5% K3Fe(CN)6 in 0.1 M NaCacodylate buffer at room temperature for 1h. After washing, cells were subsequently dehydrated through a graded ethanol series, including a bulk staining with 1% uranyl acetate at the 50% ethanol step, followed by embedding in Spurr’s resin. Ultrathin sections of a gold interference color were cut using an ultra microtome (Leica EM UC6), followed by a post-staining in a Leica EM AC20 for 40 min in uranyl acetate at 20°C and for 10 min in the lead stain at 20°C. Sections were collected on Formvar-coated copper slot grids. Grids were viewed with a JEM 1400plus transmission electron microscope (JEOL, Tokyo, Japan) operating at 80 kV.

### Confocal 3D imaging

Confocal imaging on the liver and thymic lobes was performed using the Ce3D clearing method^53^. Briefly, adult Ncr1-tdTomato thymus-grafted mice were perfused with PBS with 1% Heparin and the liver (Specifically: *lobus sinister medialis hepatis*) was fixed in 4% PFA. In addition, Ncr1-tdTomato thymic lobes from MCMV-infected neonates, controls, and NTOC lobes were fixed with PLP buffer^53^. All organs were incubated at room temperature under gentle shaking (150–220 rpm) with primary and secondary antibodies for five days each, followed by three days of Ce3D clearing solution, then mounted on glass slides and 2×2 or 3×3 tiles were imaged using a Zeiss LSM 880 AiryScan (Zaventem, Belgium) on Fast Airy mode with a Plan-Apochromat 10x/0.45, at a z-interval between 7-15 μm. Images were processed using ZEN software. Montages were made with Fiji (NIH, Bethesda, Maryland, USA), and cell quantifications in thymic lobes were performed using Arivis (Rostock, Germany). Antibodies used for 3D imaging can be found in **Table S1**.

### Multiplex ELISA

The customizable multiplex U-PLEX ELISA (MSD, Rockville, MD)) was used to measure protein concentrations of cytokines in the supernatant of NTOC culture. Murine cytokines measured: IFNγ, TNFα, GM-CSF, CCL3. NTOC supernatant was centrifuged at 10,000g to remove cells and cell debris from samples before assay measurements.

### Tissue preparation for flow cytometry and cell sorting

NTOC supernatant cells were harvested by repeated pipetting of the NTOC well and filtered after the lobes were removed from the well. Preparation of thymic lobes and spleen for single-cell suspension used for flow cytometry and cell sorting (scRNAseq analysis and OP9-DL1 co-culture) were obtained by smashing organ through a 70 μm cell strainer, the strainers were washed with 2% FCS in PBS. Perfusion was performed to remove erythrocytes from adult liver samples. The mice were anesthetized (ketamin/xylazine), and blood was extracted by retro-orbital punction followed by cardiac perfusion with 15ml of PBS with 1% heparin, using a perfusion pump. Leucocytes were extracted from adult and neonatal livers by mincing using a gentleMACS^TM^ Dissociator (Miltenyi Biotec, Cat# 130-093-235). Followed by collagenase treatment with pre-warmed RPMI containing 1 mg/mL Collagenase A (Cat# 11088793001, Merck). The resulting tissue homogenate was centrifugated on top of a 5mL 37.5% Percoll (Cat# P1644-1L, Merck) gradient (700g, 10min, 4C, break 1/acceleration 4). Adult bone marrow cells were acquired by cutting the knee-end of tibia and fibula, placing the open end downward in the tube and using centrifugal force to extract cells from bones (1 min, 1900 g, RT). Neonatal bone marrow was acquired by isolating femurs and humeri from neonates, crushing the cleaned bones with mortar and pestle, and filtering cells through a 70 μm cell strainer. To further enrich for lymphocytes, the red blood cells were lysed using ACK buffer (Lonza, Cat# 10-548E) for 2 min at RT before antibody staining. Subsequent flow cytometry or FACS prep was performed on ice. Antibody staining was performed 30min in the dark. For intracellular staining of transcription factors, cells were fixed after extracellular staining using eBioscience™ Foxp3 / Transcription Factor Staining Buffer Set (Cat# 00-5523-00, ThermoFisher). For intracellular cytokine measurements, cells were first extracted from each tissue as mentioned above and incubated with [1.5 uM] Brefeldin A (Cat# B6542-25MG, Sigma) and Monensin solution 1000x (#Cat 420701, Biolegend) for 4 h, followed by extracellular staining. The cells were then fixed and permeabilized using BD Cytofix/Cytoperm™ (Cat# 554714, BD Biosciences), and intracellular stained. Cell counts (Figure 2, 3, and 6) were derived by flow cytometry by adding Precision count beads to the sample after single cell suspension according to the manufacturer’s instructions (Biolegend, Cat# 424902), and to acquire absolute NTOC cell counts (Figure 1), they were counted on a FACSVerse using volume metrics and counting of viable and (DAPI) death cells for 15 sec immediately following single cell suspension. Fixable Viability Dye eFluor™ 780 (FVD-eF780;Cat# 65-0865-14, ThermoFisher) was used to exclude dead cells in all flow cytometry and cell sorting experiments unless otherwise stated. All antibodies used for flow cytometry and cell sorting are shown in **Table S1**. All flow cytometry antibodies were validated by supplier. Sorting of viable NTOC cells (DAPI negative) for CITE-seq and electron microscopy was performed on a BD FACSAria™ III Cell Sorter. Sorting of cell types *ex vivo* for scRNAseq (tissues) and for OP9-DL1 cultures was performed on the BD FACSymphonyTM S6 cell sorter. All flow cytometry was performed on a FACSymphony A5 Cell Analyzer. Data were analyzed with FlowJo software, version 10.8.

### Single-cell RNA seq and Cellular Indexing of Transcriptomes and Epitopes by Sequencing (CITE-seq)

**CITE-seq (Figure 1):** All viable (DAPI negative) cells from **Day 6 NTOC supernatant** (Figure 1) treated with (1) Vehicle, (2) IL-12, (3) IL-18, and (4) IL-12+IL-18 were sorted (Aria III, BD Biosciences) and pelleted by centrifugation at 400g for 5 min at 4°C. When CITE-seq was to be performed, cells were then stained with CD16/CD32 mAb (Cat# 553142, BD Bioscience) to block nonspeceific Fc receptor binding of CITE-seq antibodies for 20mins at 4°C, before being washed in excess PBS with 2% FCS used directly in downstream CITE-seq analysis. **Single-cell RNA seq (Figures 3** **& 4):** For the scRNAseq experiment comparing **tissue** group-1 ILCs from adult 8-weeks-old *Adult* and P0.5 *Neonatal* mice from *ex vivo* extracted thymus, liver, spleen, and bone marrow, the following samples were used; Adult mice: 12 thymus, 4 spleens, 12 perfused livers, and bone marrow was collected from two hind legs (tibia and fibia) of 4 mice. For all adult organs, an equal number of male and female mice were used. Neonatal mice: 90 thymus, 45 spleen, 45 bone marrow samples (arm-and leg bones from 45 neonates), 30 livers. Following single-cell suspension as described in previous paragraph, type 1 ILCs were enriched by negative selection using magnetic beads (MagniSort™ Mouse NK cell Enrichment Kit; ThermoFisher, Cat# 8804-6828-74). The samples were then sorted to enrich for viable Group-1 ILCs (CD122+Lin-: 50%), DN1-DN2 cells (CD44+Lin-: 25%), and DN3-DN4 cells (CD44-Lin-: 25%) for parallel comparison of group-1 ILCs and progenitor cells across tissues (Lin-: CD4^-^CD8a^-^CD5^-^TCRb^-^TCRgd^-^Ter-119^-^CD11c^-^F4/80^-^Ly-6G/Ly-6C^-^). Following sorting, both the NTOC supernatant cells and *ex vivo* tissue comparison cells were stained with mouse cell surface protein TotalSeq-A antibodies panels containing 9 isotype controls and 77 (NTOC) or 174 (tissue) oligo-conjugated antibodies (TotalSeq-A, Biolegend) **(Tables S2 and S3)**. The sorted single-cell suspensions were resuspended at a final estimated conc. of 1500 and 1100 cells/µl for NTOC and tissues, respectively. The cells were loaded on a Chromium GemCode Single Cell Instrument (10x Genomics) to create single-cell gel beads-in-emulsion (GEM). The scRNA-Seq libraries were prepared using GemCode Single Cell 3’ Gel Bead and Library kit, version 3 (NTOC) and version NextGEM 3.1 (tissue) (10x Genomics) according to the manufacturer’s instructions with the addition of amplification primer (3 nM, 5’CCTTGGCACCCGAGAATT*C*C) during cDNA amplification to enrich the TotalSeq-A cell surface protein oligos. Size selection with SPRIselect Reagent Kit (Beckman Coulter, B23318) was applied to separate the amplified cDNA molecules for 3’ gene expression and cell surface protein construction. TotalSeq-A protein library construction including sample index PCR using Illumina’s Truseq Small RNA primer sets and SPRIselect size selection was performed according to the manufacturer’s instructions. The cDNA content of pre-fragmentation and post-sample index PCR samples was analyzed using 2100 BioAnalyzer (Agilent). Sequencing libraries were loaded on a HiSeq4000 flow cell (NTOC) and an Illumina NovaSeq flow cell (tissue) at VIB Nucleomics core with sequencing settings according to the recommendations of 10x Genomics, pooled in a 90:10 ratio for the combined 3’ gene expression and cell surface protein samples respectively. The Cell Ranger pipeline (10x Genomics, version 3.1.0) was used to perform sample demultiplexing and to generate FASTQ files for read 1, read 2 and the i7 sample index for the gene expression and cell surface protein libraries. Read 2 of the gene expression libraries was mapped to the mouse reference genome (GRCm38.99). Subsequent barcode processing, unique molecular identifiers filtering and gene counting were performed using the Cell Ranger suite. CITE-seq reads were quantified using the feature-barcoding functionality. The average mean reads per cell across the NTOC libraries and tissue libraries were 13268 and 28039, respectively, with an average sequencing saturation of 22.9% and 51.76%, respectively, calculated by Cell Ranger (version 3.1.0). In total 4 NTOC libraries and 8 tissue libraries were created in this study, entailing 61501 and 88947 cells, respectively.

### scRNA-seq and CITE-seq processing and analysis

The aggregaton of NTOC samples was done using the Cell Ranger Aggr software from 10x Genomics. Preprocessing of the data was done by the SCRAN and Scater R packages^40^. Outlier cells were identified based on three metrics (library size, number of expressed genes and mitochondrial proportion), and cells were tagged as outliers when they were a certain median absolute deviation (MAD) away from the median value of each metric across all cells. The outliers were determined with a setting of three MADs for library size and the number of expressed genes and four MADs for mitochondrial proportion. The low-quality cells (low/high UMI counts, low/high number of genes, high % mitochondrial genes) were removed from the analysis. Genes expressed in less than 3 cells and cells expressing less than 200 genes were removed. The raw counts were processed with the Seurat R package (v 3.2.3)^54^ using the following functions with default parameters unless stated otherwise: NormalizeData, FindVariableFeatures, ScaleData, RunPCA, FindNeighbors, FindClusters (resolution: 0.8), RunUMAP (dims = 1:25). The tissue libraries were merged and processed with the Seurat package (v 4.1.1) using the following functions with default parameters unless stated otherwise: NormalizeData, FindVariableFeatues, ScaleData, RunPCA, FindNeighbors (dims = 1:30), FindClusters (resolution: 0.3), RunUMAP (dims = 1:30). Subsets were made of both datasets (NTOC dataset was filtered on ILC1s, γδT17 cells and CD4 T cells; tissue dataset was filtered on Ncr1^+^CD3e^-^ cells). These subsets were integrated and batch corrected with the Harmony package (v 0.1.0)^55^ using the following pipeline: NormalizeData, FindVariableFeatues, ScaleData, RunPCA (npcs = 150), RunHarmony, FindNeighbors (dims = 1:50), FindClusters (resolution: 1.8), RunUMAP (dims = 1:50). Clusters were further manually curated. Figures were made using the VlnPlot, DimPlot, FeaturePlot and DotPlot functions of the Seurat package.

### Statistical analysis

In all mouse experiments, the mice were randomly assigned a group. GPower3.1 software was used to predetermine sample sizes in mouse models based on mean variances of previous experiments. In case of no prior experience with the animal model a minimum of three biological replicates were sampled. The differences in variances between treatment-groups were tested with F-test, and when the F-test showed differences between conditions, statistics was performed on Log2-transformed values. In case the variances remained different on log2 transformed data a non-parametric test was used. The specific test used to evaluated differences between conditions groups are specified in the figure legend. Statistical analyses were carried out using GraphPad Prism version 9.4.1 (Graphpad Software Inc., La Jolla, CA).

## Data availability

Source data from this work has been deposited at (https://doi.org/10.6084/m9.figshare.22133480). The single cell sequencing data will be deposited in the Gene Expression Omnibus. Further information on research design is available in the Nature Research Reporting Summary.

## Extended Data Figures

**Figure S1.1:**
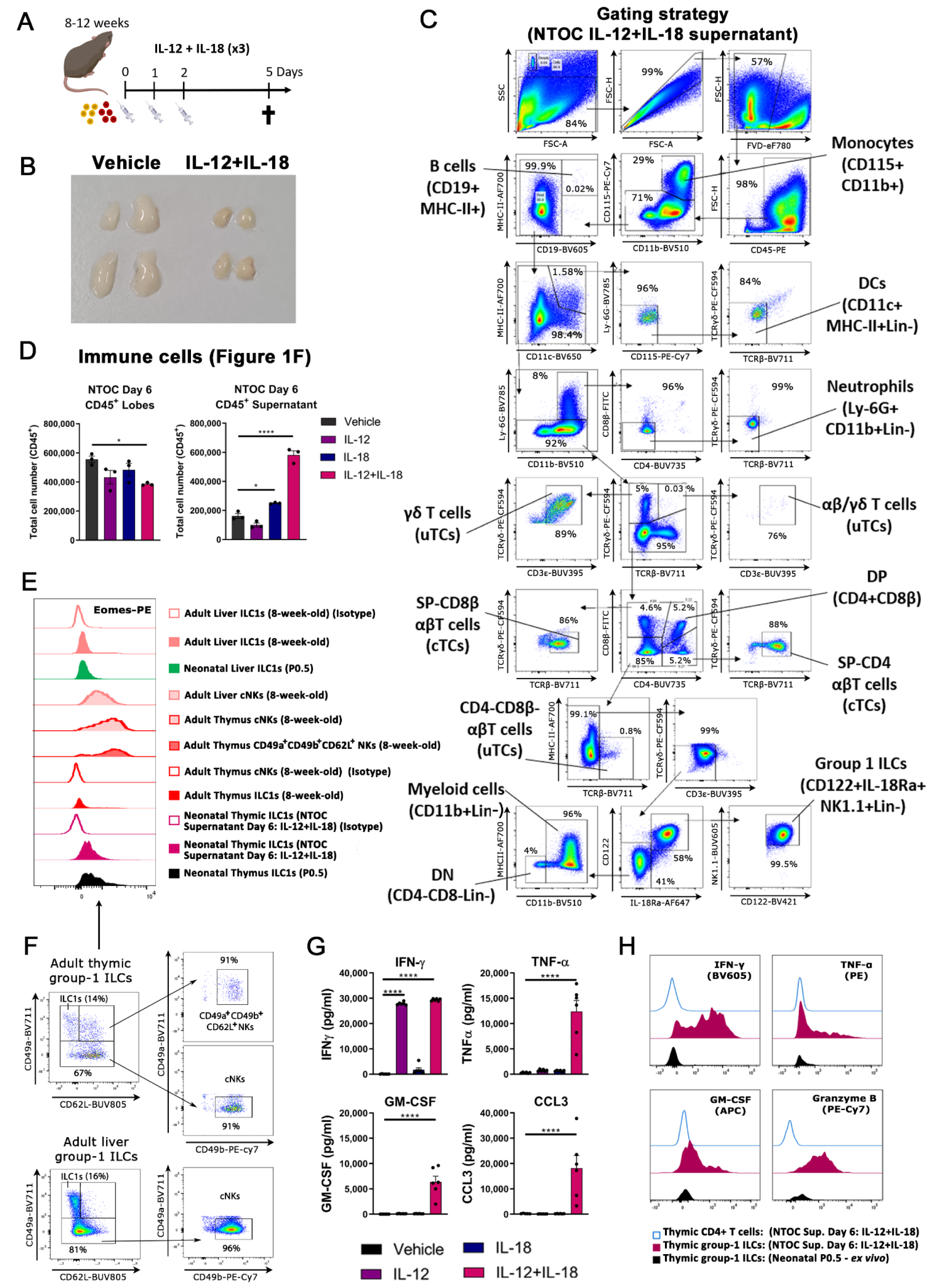
Type-1 inflammation induces expansion of thymic-exiting ILC1s from NTOC. **(A)** Schematic of cytokine injection model where adult mice are injected three times with IL12+IL-18 and terminated 5 days after the first injection. **(B)** As shown in **A**, a picture of both thymic lobes from two 8-week-old mice 5 days after the first injection with IL-12+IL-18 or Vehicle and representative of at least three independent experiments. **(C)** Flow cytometry gating strategy for figure 1E. **(D)** CD45^+^ cells in vehicle control conditions and cytokine stimulatory conditions in lobes and supernatant from three biological replicates (*n*=3) per condition. **(E)** Intracellular Eomes expression shown as representative histograms on ILC1s (CD122^+^NK1.1^+^CD49a^+^CD62L^-^Lin^-^) and cNK cells (CD122^+^NK1.1^+^CD49b^+^CD49a^-^ Lin^-^) from the liver and thymus (adults and neonates), and IL-12+IL-18 NTOC supernatant with corresponding isotype controls (open histograms), as well as unconventional adult thymic NK (CD122^+^NK1.1^+^CD49a^+^CD49b^+^CD62L^+^Lin^-^), gating shown in Figure S1F and representative of 5-6 biological replicates from two independent experiments. **(F)** Gating strategy from CD122^+^NK1.1^+^Lin^-^ gate used for differentiation of adult and neonatal ILC1s, cNK cells, and unconventional CD49a^+^CD49b^+^CD62L^+^ adult thymic NK cells in Figure S1E. **(G)** Protein measurements of cytokines in the supernatant after 6 days of NTOC, measured by multiplex ELISA from six biological replicates (*n*=6) per condition combined from three independent NTOC experiments with error bars shown as SEM. **(H)** Histograms of intracellular flow cytometry, visualizing IFNy, TNFα, GM-CSF, and Granzyme B in NTOC-derived ILC1s compared to *ex vivo* P0.5 ILC1s and internal negative control (NTOC CD4^+^ T cells) 4 hours after Brefeldin A & Monensin block and no further stimulation and representative of 3 biological replicates (*n*=3). Statistics: (**E)** One-way Anova, F test showed equal variances in treatments. Thus differences were evaluated between Vehicle and other treatments by One-way Anova (with Dunn’s multiple comparisons test), which showed significant differences between vehicle and IL-12+IL-18 lobes (effect size (170373); q-value (3.507); df (8); Adjusted p-value (0.0200)), Vehicle and IL-18 supernatant (effect size (-87923); q-value (3.419); df (8); Adjusted p-value (0.0227)), and Vehicle and IL-12+IL-18 supernatant (effect size (-419423); q-value (16.31); df (8); Adjusted p-value (<0.0001)), the enhanced thymus atrophy in NTOC IL-12+IL-18 lobes and increased supernatant cells in the IL-12+IL-18 condition is representative of at least five independent experiments. (**G)** F test showed equal variances on the log2-transformed scale for the conditions for TNF-α and CCL3. Thus differences were evaluated by One-way Anova (with Dunn’s multiple comparisons test) on log2-transformed data, which showed significant differences between treatments for TNF-α (effect size (-5.437); q-value (15.94); df (20); Adjusted p-value (<0.0001)), and for CCL3 (effect size (-6.047); q-value (12.40); df (19); Adjusted p-value (<0.0001)) – one Vehicle sample (CCL3) produced no data. F test showed unequal variances on the log2-transformed scale for the conditions for IFN-γ and GM-CSF. Thus differences were evaluated by non-parametric Kruskal-Wallis test (with Dunn’s multiple comparisons test): For IFN-γ (Kruskal-Wallis statistic (21.15), Mean rank diff. (-17.67), Z-value (4.327), Adjusted p-value (<0.0001)), and for GM-CSF (Kruskal-Wallis statistic (19.68), Mean rank diff. (-18.00), Z-value (4.409), Adjusted p-value (<0.0001)). * p < 0.05, ****p < 0.0001. **Abbreviations:** cTC = Conventional T cells; uTC = Unconventional T cells; DC = Dendritic cell; DN = Double negative (CD4 and CD8 negative thymocytes); DP = Double positive (CD4 and CD8 positive thymocytes).

**Figure S1.2:**
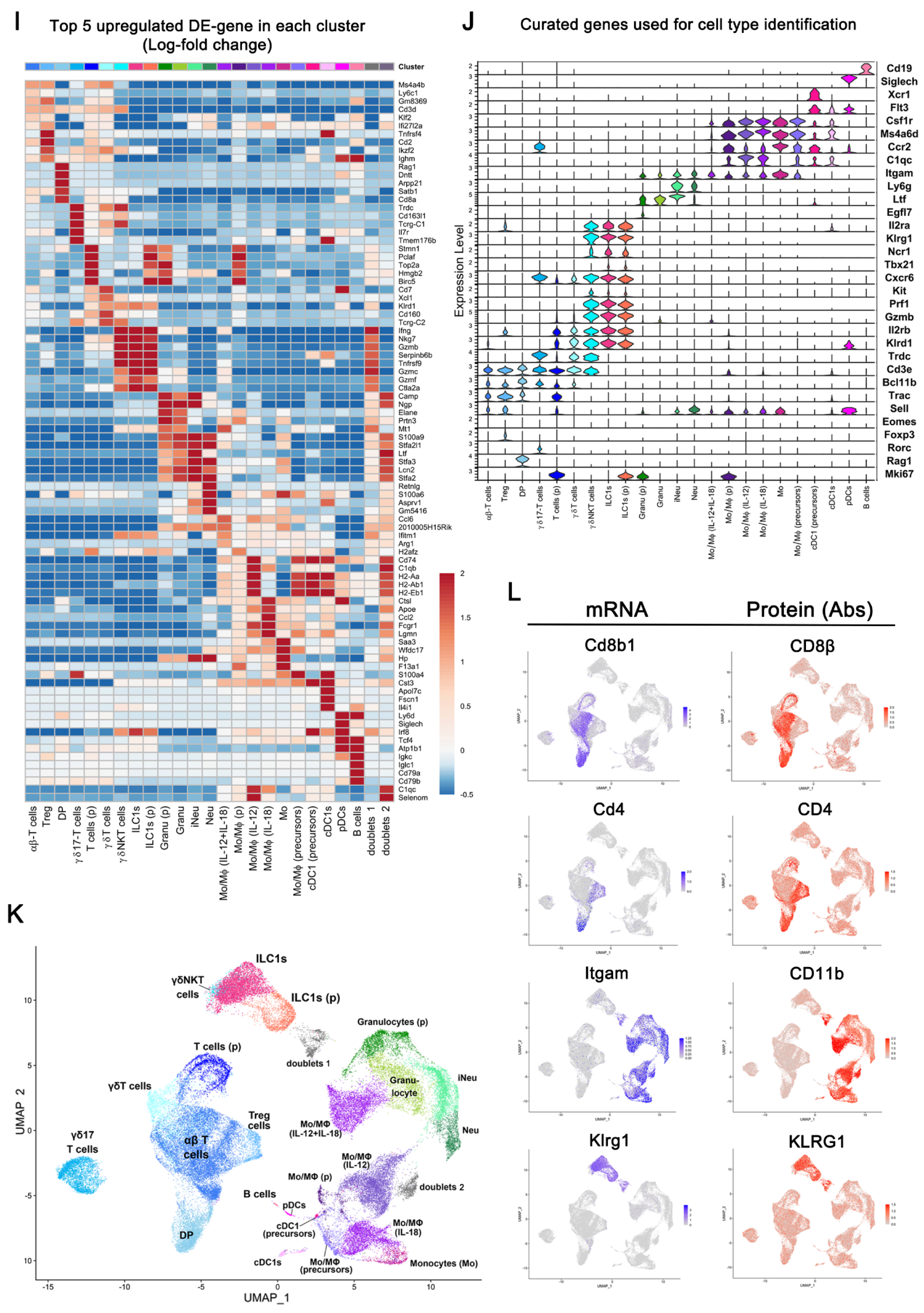
Type-1 inflammation induces expansion of thymic-exiting ILC1s from NTOC. **(I)** Heatmap showing top-5 upregulated DE-genes for each of the 23 scRNAseq clusters and doublets. **(J)** Violin plot of 23 scRNAseq clusters showing genes used for cell type and cell state identification (proliferation marker: *Mki67*). **(K)** UMAP plot of scRNAseq data, dividing the cells into 23 clusters and two clusters of doublets. **(L)** UMAP feature plots of CITE-seq data showing gene expression (mRNA) and corresponding surface protein (Abs) based on CITE-seq antibodies. **Abbreviations:** (p) = proliferating, Tc = T cell, Treg = Regulator T cell, DP = Double positive, Granu = Granunolcytes, iNeu = immature neutrophils, Neu = Neutrophils, Mo/MΦ = Monocytes/Macrophages, cDC1 = conventional type 1 DCs, pDC = plasmacytoid DCs, Bc = B cells, DE-gene = Differentially expressed gene.

**Figure S2:**
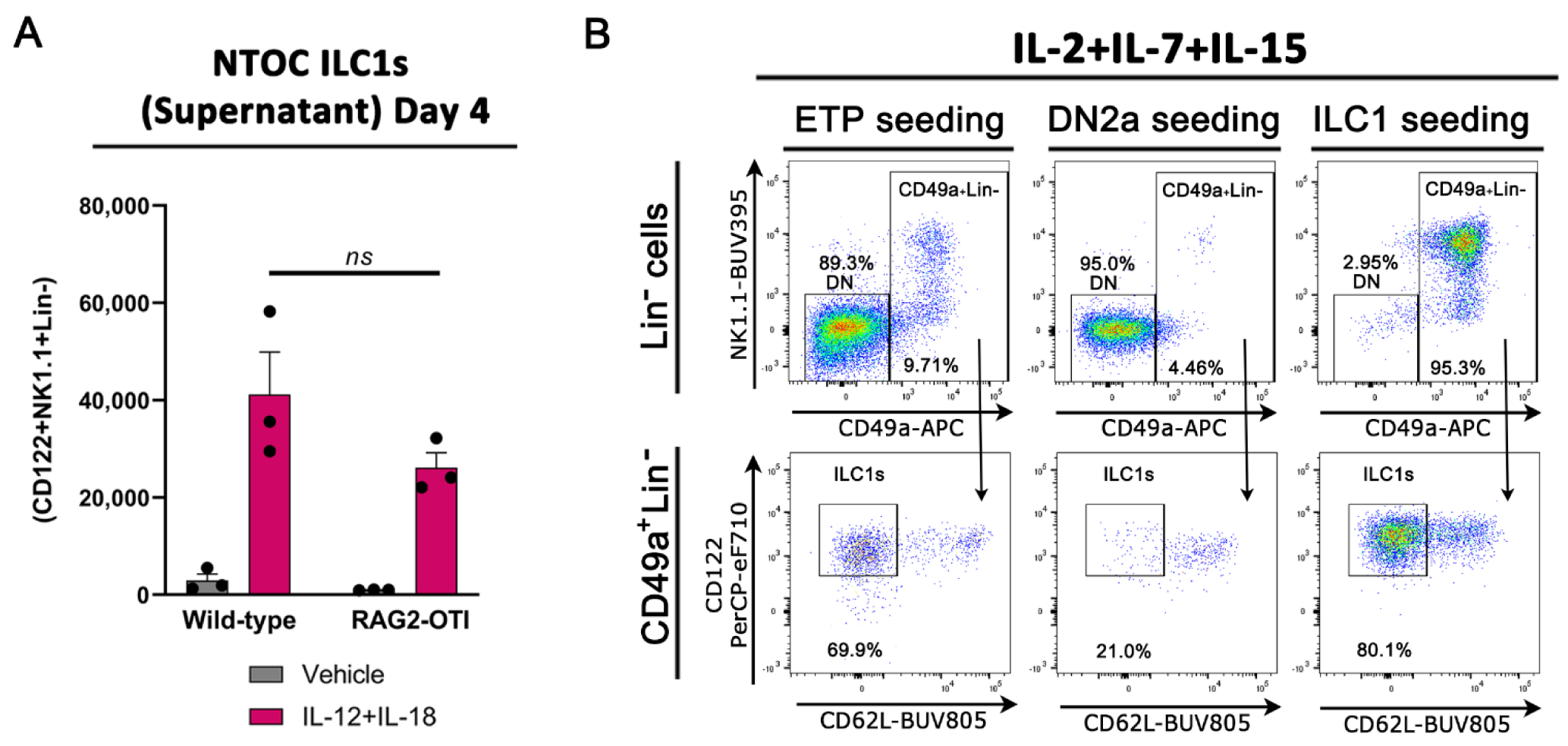
IL-12+IL-18-induced thymic ILC1s expand from immature ILC1s in a Notch-dependent manner. **(A)** ILC1s exiting NTOC into the supernatant after 4 days of culture with two lobes per condition. Thymic lobes derived from either wild-type or *Rag2*^-/-^ OTI TCR transgene mice and error bars are shown as SEM with three biological replicates (*n*=3) combined from two independent experiments. **(B)** Flow plots showing ILC1s (CD122+CD49a+CD62L-lin-) developing and expanding from (ETPs and ILC1) seeded cells following 6 days of OP9-DL1 co-culture for ETPs (CD117^(hi)^CD44^+^CD25^-^CD122^-^Lin^-^), DN2a (CD117^(hi)^CD44^+^CD25^+^CD122^-^Lin^-^), or ILC1s (CD122+Lin-) from the neonatal thymus and representative of three biological replicates (*n*=3). **(A)** Statistics; F test showed equal variances between conditions. Thus Two-way Anova was performed and found no difference between the two genotypes for the same treatment, *ns* = not significant. **Abbreviations:** DN = Double negative, ETP = Early Thymic progenitors, ns: not significant; Lin-: Lineage negative, defined as TCRβ^-^TCRγδ^-^CD3e^-^CD4^-^CD8β^-^Ly-6G^-^F4/80^-^CD19^-^CD11c^-^.

**Figure S3:**
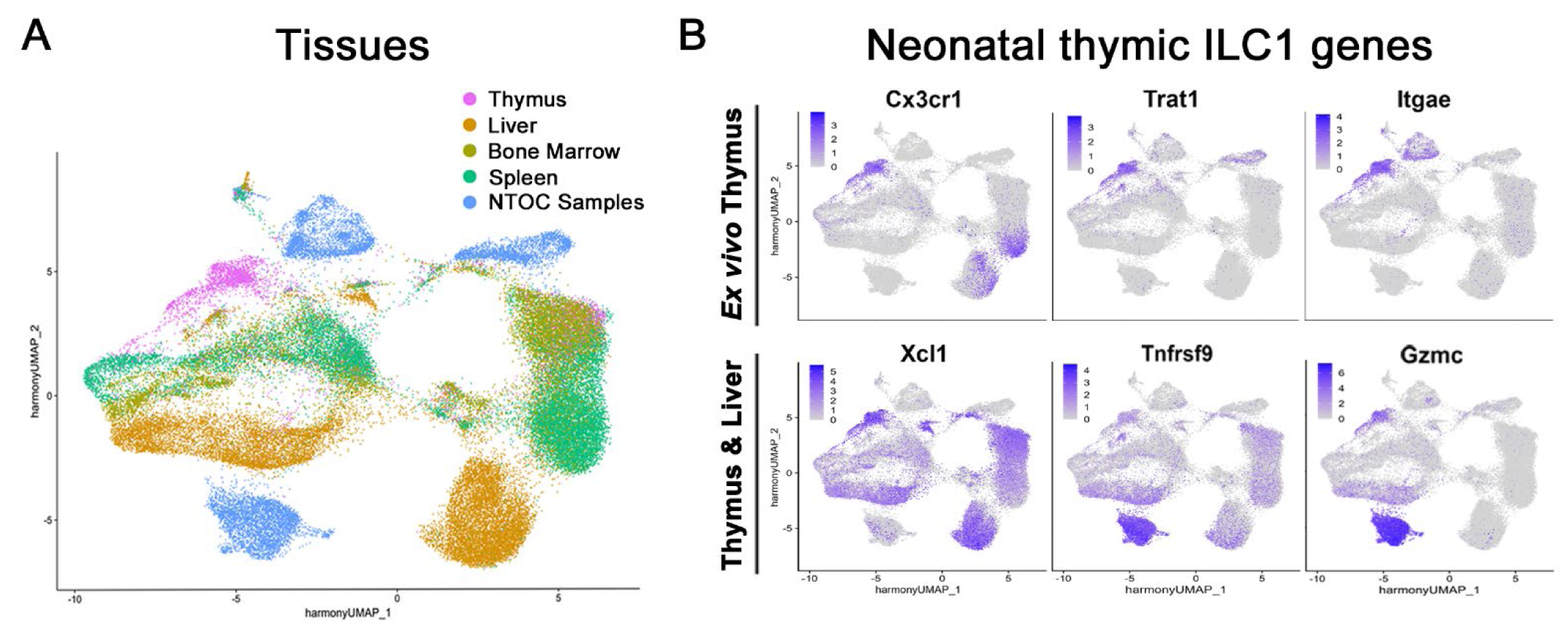
Neonatal thymic ILC1s shift expression from CX3CR1 to CXCR6 upon IL-12+IL-18 stimulation. **(A)** UMAP of Harmony integrated data from (**3A)**, showing an overview of the different tissues illustrated in **3C** (showing the neonatal and adult tissues with the same color). **(B)** UMAP feature plots of prominent curated DE-genes from neonatal thymus ILC1s.

**Figure S4:**
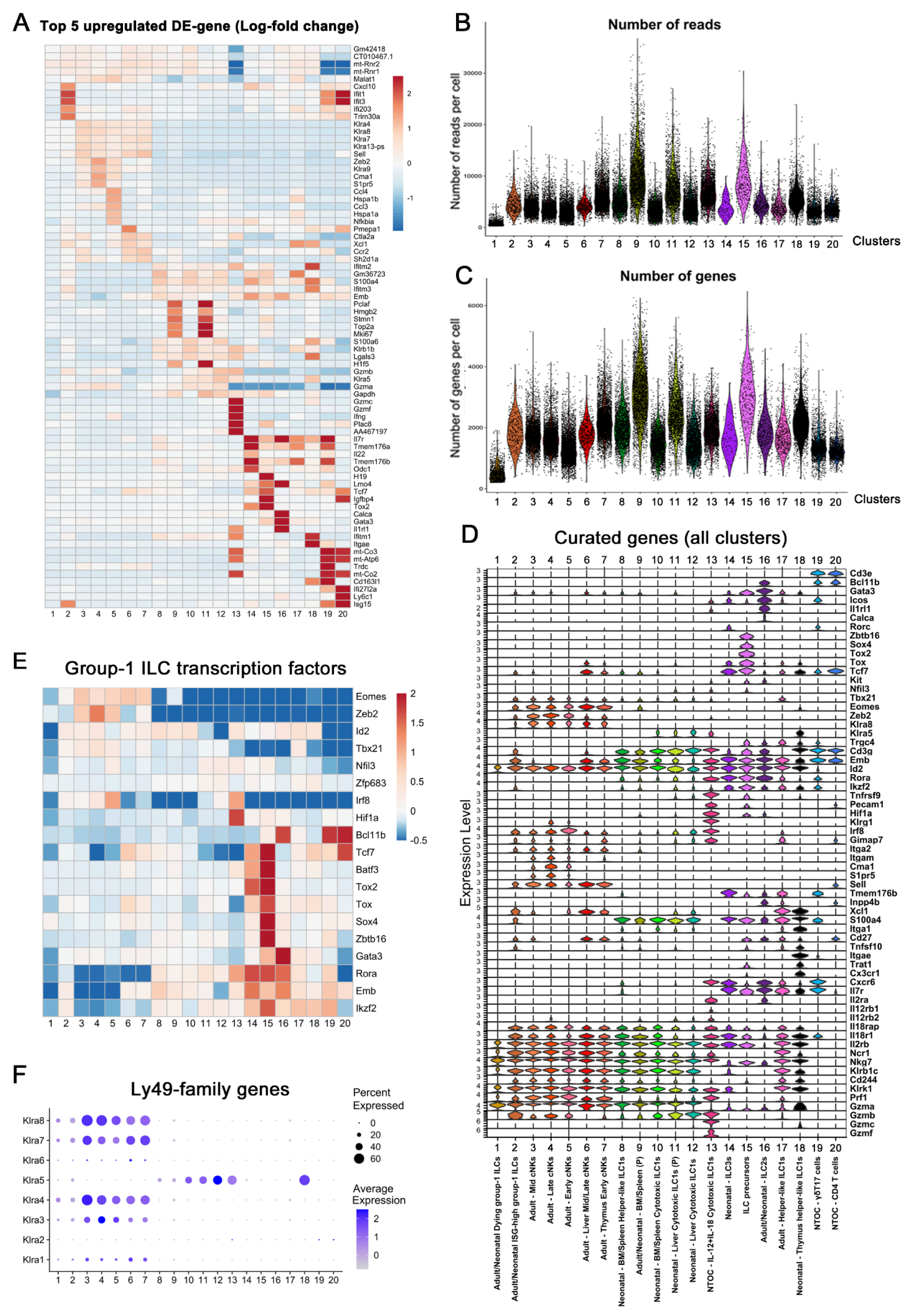
Neonatal thymic ILC1s have a unique phenotype *ex vivo*. **(A)** Top-5 upregulated DE-genes in all clusters. **(B)** Number of reads/cell for each cluster. **(C)** Number of genes/cell for each cluster. **(D)** Violin plot of gene expression in all 20 clusters with an extended list of curated genes visualizing markers for cell type, maturation, and function. **(E)** Heatmap of group-1 ILC transcription factors in all clusters. **(F)** Expression of Ly49-family genes in all clusters. **Abbreviations:** DE-gene = Differentially expressed gene.

**Figure S6:**
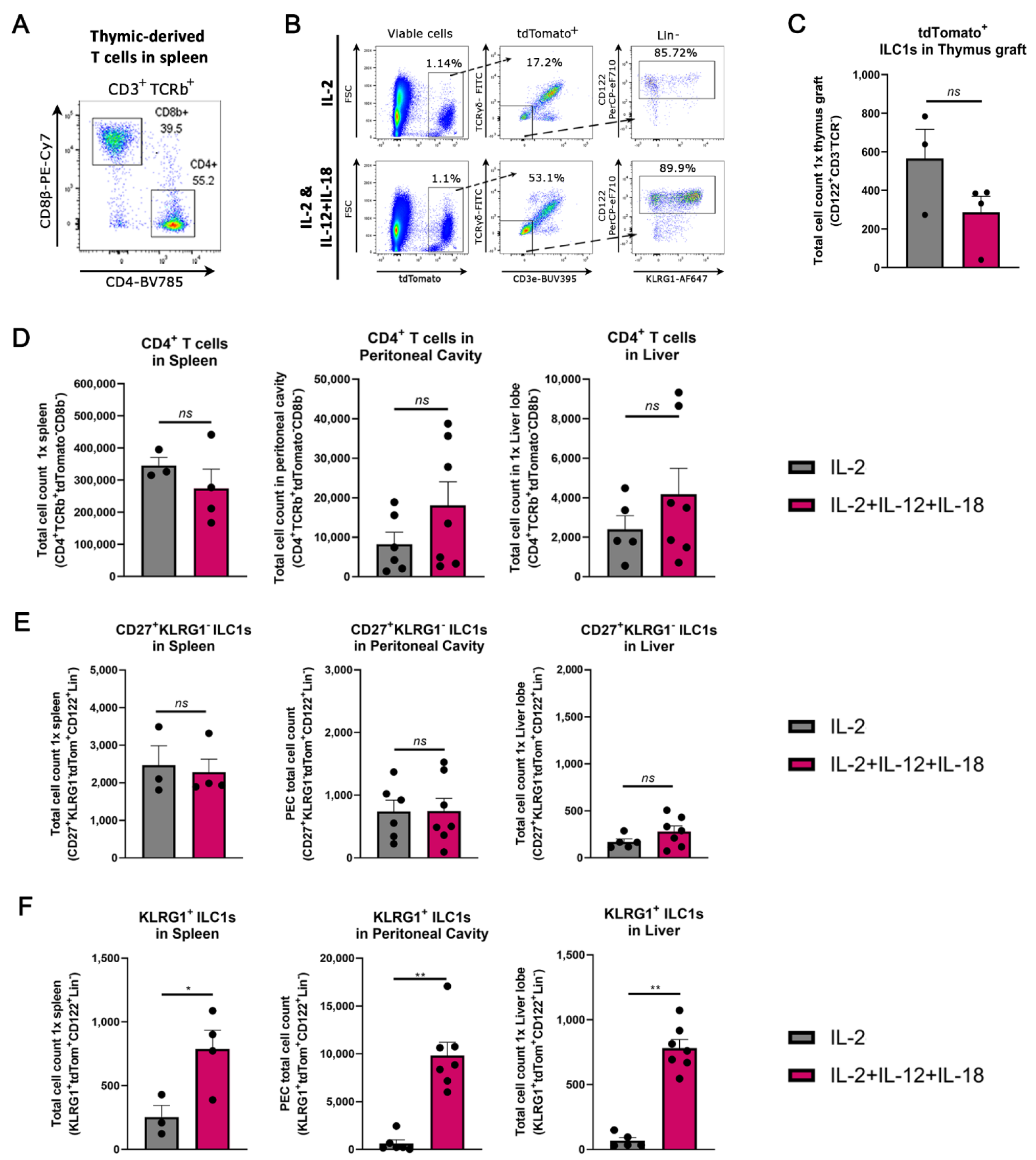
IL-12+IL-18 activated thymic ILC1s home to the liver and peritoneal cavity. **(A)** Flow plot showing splenic reconstitutions of CD4 and CD8 T cells in IL-2 injected mice 6 days after grafting representative of five biological replicates. **(B)** Gating strategy showing important gating steps for tdTomato^+^ ILC1s in IL-2 injected vs IL-2+IL-12+IL-18 injected cells from the peritoneal cavity as shown in Figure 6C. **(B)** Bar plots of tdTomato^+^ ILC1s in thymus graft. **(C)** Bar plots of thymus-graft-derived CD4 T cells in the spleen, peritoneal cavity and liver are combined from two independent experiments. **(F)** Bar plots of thymus-graft-derived CD27^+^KLRG1^-^ ILC1s in the spleen, peritoneal cavity, and liver. **(G)** Bar plots of thymus-graft-derived KLRG1^+^ ILC1s in the spleen, peritoneal cavity, and liver. Statistics; **(C-F)** F test showed unequal variances on the log2-transformed scale for the conditions for (KLRG1^+^ ILC1s in PC and Liver) Thus differences were evaluated by a non-parametric Mann-Whitney test (two-tailed), KLRG1^+^ ILC1s in PC (effect size (8614); p-value (0.0012)) and KLRG1^+^ ILC1s in Liver (effect size (725); p-value (0.0025)). All other conditions were evaluated by a T test (two-tailed), KLRG1^+^ ILC1s in spleen (effect size (787.5); p-value (0.0380)): * p< 0.05, ** p < 0.01, *ns* = Not significant. **Abbreviations:** Lin-: Lineage negative, defined as: TCRβ^-^TCRγδ^-^CD3e^-^CD4^-^ CD8β^-^Ter-119^-^Ly-6G^-^.

## Supplementary Information

**Supplementary Table 1:** Flow cytometry and cell sorting antibodies.

| Antigen | Fluorochrome | Clone | Source | Cat# | Dilution (1:X) | Notes |
| --- | --- | --- | --- | --- | --- | --- |
| CD49b | FITC | DX5 | ThermoFisher | 11-5971 | 1/200 |  |
| CD122 | BV421 | TM-β1 | BD Biosciences | 752988 | 1/50 |  |
| CD49a | BV711 | Ha31/8 | BD Biosciences | 564863 | 1/100 |  |
| NK-1.1 | BUV615 | PK136 | BD Biosciences | 751111 | 1/100 |  |
| CD62L | BUV805 | MEL-14 | BD Biosciences | 741924 | 1/400 |  |
| CD27 | BUV737 | LG.3A10 | BD Biosciences | 612831 | 1/100 |  |
| KLRG1 | PE-Cy5 | 2F1 | ThermoFisher | 15-5893-80 | 1/400 |  |
| CD8β | BV510 | YTS156.7.7 | BioLegend | 126631 | 1/400 | For Lineage |
| Ter-119 | BV785 | TER-119 | BioLegend | 116245 | 1/200 | For Lineage |
| Ly-6G | BV785 | 1A8 | BioLegend | 127645 | 1/200 | For Lineage |
| F4-80 | BV785 | BM8 | BioLegend | 123141 | 1/200 | For Lineage |
| CD11c | BV785 | N418 | BioLegend | 117336 | 1/200 | For Lineage |
| CD19 | BV785 | 6D5 | BioLegend | 115543 | 1/100 | For Lineage |
| CD19 | AF700 | 1D3 | ThermoFisher | 56-0193-82 | 1/100 | For Lineage |
| TCRγδ | PE-CF594 | GL3 | BD Biosciences | 563532 | 1/200 | For Lineage |
| TCRβ | PE-CF594 | H57-597 | BD Biosciences | 562841 | 1/100 | For Lineage |
| CD3e | BUV395 | 145-2C11 | BD Bioscience | 563565 | 1/100 | For Lineage |
| CD4 | BUV615 | RM4-5 | BD Biosciences | 751486 | 1/200 | For Lineage |
| CD4 | BUV737 | GK1.5 | BD Bioscience | 612761 | 1/200 | For Lineage |
| B220 | AF700 | RA3-6B2 | BioLegend | 103232 | 1/200 |  |
| CXCR6 | PE | SA051D1 | BioLegend | 151103 | 1/100 |  |
| CX3CR1 | FITC | SA011F11 | BioLegend | 149020 | 1/50 |  |
| CD127 | BV605 | A7R34 | BioLegend | 135041 | 1/50 |  |
| CD103 | BUV395 | M290 | BD Bioscience | 740238 | 1/100 |  |
| CD49b | PE-Cy7 | DX5 | BioLegend | 108921 | 1/100 |  |
| Eomes | PE | Dan11mag | ThermoFisher | 12-4875-82 | 1/100 |  |
| Rat IgG2a-k Isotype Control | PE | eBR2a | ThermoFisher | 12-4321-42 | 1/100 | Eomes Isotype control |
| Eomes | AF647 | W17001A | Biolegend | 157703 | 1/100 | Intracellular |
| CD31 (PECAM.1) | FITC | 390 | ThermoFisher | 11-0311-82 | 1/100 |  |
| CD200R | PE | OX-110 | BioLegend | 123907 | 1/100 |  |
| Ki-67 | BV605 | 16A8 | BioLegend | 652413 | 1/50 | Intracellular |
| T-bet | PE-Cy7 | 4B10 | ThermoFisher | 25-5825-82 | 1/50 | Intracellular |
| IFN-γ | BV605 | XMG1.2 | Biolegend | 505839 | 1/100 | Intracellular |
| GM-CSF | APC | MP1-22E9 | ThermoFisher | 17-7331-82 | 1/100 | Intracellular |
| TNF-α | PE | MP6-XT22 | ThermoFisher | 12-7321-81 | 1/100 | Intracellular |
| Granzyme B | PE-Cy7 | NGZB | ThermoFisher | 25-8898-80 | 1/100 | Intracellular |
| Collagen IV | Rabbit IgG (non-conjugated) | Polyclonal | Abcam | ab19808 | 1/100 | Used in 3D imaging |
| Secondary Rabbit-IgG | Dylight488 | Polyclonal | ThermoFisher | SA5-10038 | 1/500 | Used in 3D imaging |
| CD3e | AF647 | 17A2 | Biolegend | 100209 | 1/100 | Used in 3D imaging |
| CD16/32 | purified | 2.4G2 | BD Biosciences | 553142 | 1/100 | For Fc-Receptor blocking |
| Ly-6G | FITC | 1A8 | BD Bioscience | 551460 | 1/100 |  |
| KLRG1 | BV605 | 2F1 | BioLegend | 138419 | 1/400 |  |
| KLRG1 | AF561 | 2F1 | ThermoFisher | 505-5893-82 | 1/400 |  |
| CD11c | BV650 | N418 | BioLegend | 117339 | 1/100 |  |
| TCRβ | BV711 | H57-597 | BioLegend | 109243 | 1/100 | |
| MHC-II | BV786 | 2G9 | BD Biosciences | 743875 | 1/800 |  |
| IL-18Ra | AF647 | BG/IL18RA | Biolegend | 132903 | 1/100 |  |
| CD45 | PE | 30-F11 | BioLegend | 103105 | 1/300 |  |
| CD11b | PE-Cy5 | M1/70 | ThermoFisher | 15-0112-81 | 1/800 |  |
| CD115 | PE-Cy7 | AFS98 | BioLegend | 135523 | 1/100 |  |
| CD115 | BV650 | AFS98 | BD Biosciences | 750891 | 1/100 |  |
| MHC-II | AF700 | M5/114.15.2 | BioLegend | 107622 | 1/600 |  |
| CD19 | BV605 | 6D5 | BioLegend | 115540 | 1/200 |  |
| CD11b | BV510 | M1/70 | BD Biosciences | 562950 | 1/600 |  |
| NKp46 | BUV737 | 29A1.4 | BD Bioscience | 612805 | 1/50 |  |
| CD117 | BB515 | 2B8 | BD Biosciences | 564481 | 1/100 | Not available from supplier anymore |
| CD117 | FITC | 2B8 | BD Biosciences | 564481 | 1/100 |  |
| CD127 | BV421 | A7R34 | BioLegend | 135027 | 1/100 |  |
| CD25 | BV605 | PC61 | BD Bioscience | 563061 | 1/400 |  |
| CD4 | APC | GK1.5 | BioLegend | 100426 | 1/200 |  |
| CD44 | PE | IM7 | BD Biosciences | 553134 | 1/400 |  |
| CD28 | PE-Cy7 | E18 | Biolegend | 122014 | 1/100 |  |
| KLRG1 | AF647 | 2F1 | ThermoFisher | 51-5893-82 | 1/200 |  |
| NK-1.1 | PE-Cy7 | PK136 | BD biosciences | 552878 | 1/100 |  |
| CD4 | BV510 | RM4-5 | Biolegend | 100559 | 1/200 |  |
| NK-1.1 | APC | PK136 | BioLegend | 108710 | 1/100 |  |
| CD4 | FITC | GK1.5 | BD Biosciences | 557307 | 1/200 |  |
| CD122 | PerCP-eFluor710 | TM-b1 (TM-beta1) | ThermoFisher | 46-1222-82 | 1/100 |  |
| CD8β | PE-Cy7 | YTS156.7.7 | Biolegend | 126616 | 1/200 | |
| CD19 | APC | 1D3 | BioLegend | 152410 | 1/200 |  |
| **CD11c** | **APC** | **HL3** | **BD Biosciences** | **550261** | **1/200** |  |
| **CD11b** | **BV605** | **M1/70** | **BD Horizon** | **563015** | **1/400** |  |
| **MHC-II** | **PE** | **M5/114.15.2** | **ThermoFisher** | **12-5321-81** | **1/600** |  |

**Supplementary Table 2:** Total-seq antibodies (NTOC)

| **TotalSeq** | **TotalSeq-A ID** | **Target** | **Clone** | **Isotype** |
| --- | --- | --- | --- | --- |
| A | 0001 | CD4 | RM4-5 | Rat IgG2a, κ |
| A | 0002 | CD8a | 53-6.7 | Rat IgG2a, κ |
| A | 0003 | CD366 | RMT3-23 | Rat IgG2a, κ |
| A | 0004 | CD279 | RMP1-30 | Rat IgG2b, κ |
| A | 0012 | CD117 | 2B8 | Rat IgG2b, κ |
| A | 0013 | Ly-6C | HK1.4 | Rat IgG2c, κ |
| A | 0014 | CD11b | M1/70 | Rat IgG2b, κ |
| A | 0015 | Ly-6G | 1A8 | Rat IgG2a, κ |
| A | 0070 | CD49f | GoH3 | Rat IgG2a, κ |
| A | 0073 | CD44 | IM7 | Rat IgG2a, κ |
| A | 0074 | CD54 | YN1/1.7.4 | Rat IgG2b, κ |
| A | 0076 | CD15 | MC-480 | Mouse IgM, κ |
| A | 0077 | CD73 | TY/11.8 | Rat IgG1, κ |
| A | 0078 | CD49d | R1-2 | Rat IgG2b, κ |
| A | 0079 | CD200 | OX-90 | Rat IgG2a, κ |
| A | 0090 | IgG1, κ Isotype Ctrl | MOPC-21 | Mouse<br>(BALB/c)<br>IgG1, α<sub>1</sub> |
| A | 0091 | IgG2a, κ Isotype Ctrl | MOPC-173 | Mouse IgG2a,<br>α<sub>1</sub> |
| A | 0092 | IgG2b, κ Isotype Ctrl | MPC-11 | Mouse IgG2b,<br>α<sub>1</sub> |
| A | 0095 | IgG2b, κ Isotype Ctrl | RTK4530 | Rat IgG2b, α<sub>1</sub> |
| A | 0097 | CD25 | PC61 | Rat IgG1, λ |
| A | 0098 | CD135 | A2F10 | Rat IgG2a, κ |
| A | 0104 | CD102 | 3C4 (MIC2/4) | Rat IgG2a, κ |
| A | 0106 | CD11c | N418 | Armenian<br>Hamster IgG |
| A | 0107 | CD21,CD35 | 7E9 | Rat IgG2a, κ |
| A | 0108 | CD23 | B3B4 | Rat IgG2a, κ |
| A | 0110 | CD43 | S11 | Rat IgG2b |
| A | 0112 | CD62L | MEL-14 | Rat IgG2a, κ |
| A | 0114 | F4/80 | BM8 | Rat IgG2a, κ |
| A | 0115 | FcεRIα | MAR-1 | Armenian<br>Hamster IgG |
| A | 0117 | I-A/I-E | M5/114.15.2 | Rat IgG2b, κ |
| A | 0118 | NK-1.1 | PK136 | Mouse IgG2a,<br>κ |
| A | 0119 | Siglec H | 551 | Rat IgG1, κ |
| A | 0120 | TCR β chain | H57-597 | Armenian<br>Hamster IgG |
| A | 0121 | TCR γ/δ | GL3 | Armenian<br>Hamster IgG |
| A | 0122 | TER-119 | TER-119 | Rat IgG2b, κ |
| A | 0130 | Ly-6A/E | D7 | Rat IgG2a, κ |
| A | 0171 | CD278 | C398.4A | Armenian<br>Hamster IgG |
| A | 0184 | CD335 | 29A1.4 | Rat IgG2a, κ |
| A | 0190 | CD274 | MIH6 | Rat IgG2a, κ |
| A | 0191 | CD27 | LG.3A10 | Armenian<br>Hamster IgG |
| A | 0192 | CD20 | SA275A11 | Rat IgG2b, κ |
| A | 0193 | CD357 | DTA-1 | Rat IgG2b, λ |
| A | 0194 | CD137 | 17B5 | Syrian<br>Hamster IgG |
| A | 0195 | CD134 | OX-86 | Rat IgG1, κ |
| A | 0197 | CD69 | H1.2F3 | Armenian<br>Hamster IgG |
| A | 0198 | CD127 | A7R34 | Rat IgG2a, κ |
| A | 0200 | CD86 | GL-1 | Rat IgG2a, κ |
| A | 0201 | CD103 | 2E7 | Armenian<br>Hamster IgG |
| A | 0203 | CD150 | TC15-12F12.2 | Rat IgG2a, λ |
| A | 0204 | CD28 | 37.51 | Syrian<br>Hamster IgG |
| A | 0209 | TCR Vγ1.1 | 2.11 | Armenian<br>Hamster IgG |
| A | 0210 | TCR Vγ3 | 536 | Syrian<br>Hamster IgG |
| A | 0211 | TCR Vγ2 | UC3-10A6 | Armenian<br>Hamster IgG |
| A | 0212 | CD24 | M1/69 | Rat IgG2b, $\kappa$ |
| A | 0214 | Integrin β7 | FIB504 | Rat IgG2a, $\kappa$ |
| A | 0225 | CD196 | 29-2L17 | Armenian<br>Hamster IgG |
| A | 0226 | CD106 | 429 (MVCAM.A) | Rat IgG2a, $\kappa$ |
| A | 0227 | CD122 | 5H4 | Rat IgG2a, $\kappa$ |
| A | 0228 | CD183 | CXCR3-173 | Armenian<br>Hamster IgG |
| A | 0229 | CD62P | RMP-1 | Mouse IgG2a,<br>$\kappa$ |
| A | 0230 | CD8b | YTS156.7.7 | Rat IgG2b, $\kappa$ |
| A | 0232 | MAdCAM-1 | MECA-367 | Rat IgG2a, $\kappa$ |
| A | 0235 | TCR Vβ8.1,8.2 | KJ16-133.18 | Rat IgG2a, $\kappa$ |
| A | 0236 | IgG1, $\kappa$ Isotype Ctrl | RTK2071 | Rat IgG1, $\Theta$] |
| A | 0237 | IgG1, $\lambda$ Isotype Ctrl | G0114F7 | Rat IgG1, $\Theta^a$ |
| A | 0238 | IgG2a, $\kappa$ Isotype Ctrl | RTK2758 | Rat IgG2a, $\Theta$] |
| A | 0240 | Rat IgG2c, $\kappa$ Isotype<br>Ctrl | RTK4174 | Rat IgG2c, $\Theta$] |
| A | 0241 | IgG Isotype Ctrl | HTK888 | Armenian<br>Hamster IgG |
| A | 0250 | KLRG1 | 2F1/KLRG1 | Syrian<br>Hamster IgG |
| A | 0354 | TCR Vβ5.1, 5.2 | MR9-4 | Mouse IgG1,<br>$\kappa$ |
| A | 0376 | CD195 | HM-CCR5 | Armenian<br>Hamster IgG |
| A | 0377 | CD197 | 4B12 | Rat IgG2a, $\kappa$ |
| A | 0378 | CD223 | C9B7W | Rat IgG1, $\kappa$ |
| A | 0379 | CD62E | RME-1/CD62E | Mouse IgG1,<br>$\kappa$ |
| A | 0381 | Panendothelial Cell<br>Antigen | MECA-32 | Rat IgG2a, $\kappa$ |
| A | 0388 | CD152 | UC10-4B9 | Armenian<br>Hamster IgG |
| A | 0554 | CD309 | 89B3A5 | Rat IgG2a, $\kappa$ |

**Supplementary Table 3:** Total-seq antibodies (Tissues)

| TotalSeq | TotalSeq-A ID | Target | Clone | Isotype |
| --- | --- | --- | --- | --- |
| A | 0001 | CD4 | RM4-5 | Rat IgG2a, κ |
| A | 0002 | CD8a | 53-6.7 | Rat IgG2a, κ |
| A | 0003 | CD366 | RMT3-23 | Rat IgG2a, κ |
| A | 0004 | CD279 | RMP1-30 | Rat IgG2b, κ |
| A | 0012 | CD117 | 2B8 | Rat IgG2b, κ |
| A | 0013 | Ly-6C | HK1.4 | Rat IgG2c, κ |
| A | 0014 | CD11b | M1/70 | Rat IgG2b, κ |
| A | 0015 | Ly-6G | 1A8 | Rat IgG2a, κ |
| A | 0070 | CD49f | GoH3 | Rat IgG2a, κ |
| A | 0074 | CD54 | YN1/1.7.4 | Rat IgG2b, κ |
| A | 0075 | CD90.2 | 30-H12 | Rat IgG2b, κ |
| A | 0076 | CD15 | MC-480 | Mouse IgM, κ |
| A | 0077 | CD73 | TY/11.8 | Rat IgG1, κ |
| A | 0078 | CD49d | R1-2 | Rat IgG2b, κ |
| A | 0079 | CD200 | OX-90 | Rat IgG2a, κ |
| A | 0090 | IgG1, κ Isotype Ctrl | MOPC-21 | Mouse (BALB/c) IgG1, ♂ |
| A | 0091 | IgG2a, κ Isotype Ctrl | MOPC-173 | Mouse IgG2a, ♂ |
| A | 0092 | IgG2b, κ Isotype Ctrl | MPC-11 | Mouse IgG2b, ♂ |
| A | 0093 | CD19 | 6D5 | Rat IgG2a, κ |
| A | 0095 | IgG2b, κ Isotype Ctrl | RTK4530 | Rat IgG2b, ♂ |
| A | 0097 | CD25 | PC61 | Rat IgG1, λ |
| A | 0098 | CD135 | A2F10 | Rat IgG2a, κ |
| A | 0103 | CD45R/B220 | RA3-6B2 | Rat IgG2a, κ |
| A | 0104 | CD102 | 3C4 (MIC2/4) | Rat IgG2a, κ |
| A | 0105 | CD115 | AFS98 | Rat IgG2a, κ |
| A | 0106 | CD11c | N418 | Armenian Hamster IgG |
| A | 0107 | CD21,CD35 | 7E9 | Rat IgG2a, κ |
| A | 0108 | CD23 | B3B4 | Rat IgG2a, κ |
| A | 0109 | CD16/32 | 93 | Rat IgG2a, λ |
| A | 0110 | CD43 | S11 | Rat IgG2b |
| A | 0111 | CD5 | 53-7.3 | Rat IgG2a, κ |
| A | 0112 | CD62L | MEL-14 | Rat IgG2a, κ |
| A | 0113 | CD93 | AA4.1 | Rat IgG2b, κ |
| A | 0114 | F4/80 | BM8 | Rat IgG2a, κ |
| A | 0115 | FcεRIα | MAR-1 | Armenian<br>Hamster IgG |
| A | 0117 | I-A/I-E | M5/114.15.2 | Rat IgG2b, κ |
| A | 0118 | NK-1.1 | PK136 | Mouse IgG2a,<br>κ |
| A | 0119 | Siglec H | 551 | Rat IgG1, κ |
| A | 0120 | TCR β chain | H57-597 | Armenian<br>Hamster IgG |
| A | 0121 | TCR γ/δ | GL3 | Armenian<br>Hamster IgG |
| A | 0122 | TER-119 | TER-119 | Rat IgG2b, κ |
| A | 0130 | Ly-6A/E | D7 | Rat IgG2a, κ |
| A | 0134 | CD146 | P1H12 | Mouse IgG1,<br>κ |
| A | 0171 | CD278 | C398.4A | Armenian<br>Hamster IgG |
| A | 0173 | CD206 | C068C2 | Rat IgG2a, κ |
| A | 0182 | CD3 | 17A2 | Rat IgG2b, κ |
| A | 0184 | CD335 | 29A1.4 | Rat IgG2a, κ |
| A | 0190 | CD274 | MIH6 | Rat IgG2a, κ |
| A | 0191 | CD27 | LG.3A10 | Armenian<br>Hamster IgG |
| A | 0192 | CD20 | SA275A11 | Rat IgG2b, κ |
| A | 0193 | CD357 | DTA-1 | Rat IgG2b, λ |
| A | 0194 | CD137 | 17B5 | Syrian<br>Hamster IgG |
| A | 0195 | CD134 | OX-86 | Rat IgG1, κ |
| A | 0197 | CD69 | H1.2F3 | Armenian<br>Hamster IgG |
| A | 0198 | CD127 | A7R34 | Rat IgG2a, κ |
| A | 0200 | CD86 | GL-1 | Rat IgG2a, κ |
| A | 0201 | CD103 | 2E7 | Armenian<br>Hamster IgG |
| A | 0202 | CD64 | X54-5/7.1 | Mouse IgG1, κ |
| A | 0203 | CD150 | TC15-12F12.2 | Rat IgG2a, λ |
| A | 0204 | CD28 | 37.51 | Syrian Hamster IgG |
| A | 0209 | TCR Vγ1.1 | 2.11 | Armenian Hamster IgG |
| A | 0210 | TCR Vγ3 | 536 | Syrian Hamster IgG |
| A | 0211 | TCR Vγ2 | UC3-10A6 | Armenian Hamster IgG |
| A | 0212 | CD24 | M1/69 | Rat IgG2b, κ |
| A | 0214 | Integrin β7 | FIB504 | Rat IgG2a, κ |
| A | 0222 | ERK1 | W15133A | Rat IgG2a, κ |
| A | 0223 | RORγ | 2F7-2 | Mouse IgG2a, κ |
| A | 0225 | CD196 | 29-2L17 | Armenian Hamster IgG |
| A | 0226 | CD106 | 429 (MVCAM.A) | Rat IgG2a, κ |
| A | 0227 | CD122 | 5H4 | Rat IgG2a, κ |
| A | 0228 | CD183 | CXCR3-173 | Armenian Hamster IgG |
| A | 0229 | CD62P | RMP-1 | Mouse IgG2a, κ |
| A | 0230 | CD8b | YTS156.7.7 | Rat IgG2b, κ |
| A | 0232 | MAdCAM-1 | MECA-367 | Rat IgG2a, κ |
| A | 0235 | TCR Vβ8.1,8.2 | KJ16-133.18 | Rat IgG2a, κ |
| A | 0236 | IgG1, κ Isotype Ctrl | RTK2071 | Rat IgG1, ♂ |
| A | 0237 | IgG1, λ Isotype Ctrl | G0114F7 | Rat IgG1, ♂<sup>a</sup> |
| A | 0238 | IgG2a, κ Isotype Ctrl | RTK2758 | Rat IgG2a, ♂ |
| A | 0239 | IgG2a | #N/A | #N/A |
| A | 0240 | Rat IgG2c, κ Isotype Ctrl | RTK4174 | Rat IgG2c, ♂ |
| A | 0241 | IgG Isotype Ctrl | HTK888 | Armenian Hamster IgG |
| A | 0249 | IRF4 | IRF4.3E4 | Rat IgG1, κ |
| A | 0250 | KLRG1 | 2F1/KLRG1 | Syrian Hamster IgG |
| A | 0354 | TCR Vβ5.1, 5.2 | MR9-4 | Mouse IgG1,<br>κ |
| A | 0376 | CD195 | HM-CCR5 | Armenian<br>Hamster IgG |
| A | 0377 | CD197 | 4B12 | Rat IgG2a, κ |
| A | 0378 | CD223 | C9B7W | Rat IgG1, κ |
| A | 0379 | CD62E | RME-1/CD62E | Mouse IgG1,<br>κ |
| A | 0380 | CD90/CD90.1 | OX-7 | #N/A |
| A | 0381 | Panendothelial Cell<br>Antigen | MECA-32 | Rat IgG2a, κ |
| A | 0388 | CD152 | UC10-4B9 | Armenian<br>Hamster IgG |
| A | 0415 | P2RY12 | S16007D | Rat IgG2b, κ |
| A | 0416 | CD300LG | ZAQ5 | Rat IgG2a, κ |
| A | 0417 | CD163 | S15049I | Rat IgG2a, κ |
| A | 0421 | CD49b | HMa2 | Armenian<br>Hamster IgG |
| A | 0422 | CD172a (SIRPα) | P84 | Rat IgG1, κ |
| A | 0424 | CD14 | Sa14-2 | Rat IgG2a, κ |
| A | 0426 | CD192 (CCR2) | SA203G11 | Rat IgG2b, κ |
| A | 0429 | CD48 | HM48-1 | Armenian<br>Hamster IgG |
| A | 0434 | ReceptorD4 | #N/A | #N/A |
| A | 0435 | GABRB3 | #N/A | #N/A |
| A | 0437 | CD207_mh | 4C7 | Mouse IgG2a,<br>κ |
| A | 0438 | KCC2 | #N/A | #N/A |
| A | 0439 | CD201 | RCR-16 | Rat IgG2a, κ |
| A | 0440 | CD169 | 3D6.112 | Rat IgG2a, κ |
| A | 0441 | CD71 | RI7217 | Rat IgG2a, κ |
| A | 0442 | Notch 1 | HMN1-12 | Armenian<br>Hamster IgG |
| A | 0443 | CD41 | MWReg30 | Rat IgG1, κ |
| A | 0444 | CD184 | L276F12 | Rat IgG2b, κ |
| A | 0448 | CD204 | 1F8C33 | Rat IgG2a |
| A | 0449 | CD326 | G8.8 | Rat IgG2a, κ |
| A | 0450 | IgM | RMM-1 | Rat IgG2a, κ |
| A | 0551 | CD301a | LOM-8.7 | Rat IgG2a, κ |
| A | 0552 | CD304 | 3E12 | Rat IgG2a, κ |
| A | 0554 | CD309 | 89B3A5 | Rat IgG2a, κ |
| A | 0555 | CD36 | HM36 | Armenian<br>Hamster IgG |
| A | 0556 | CD370 | 7H11 | Rat IgG1, κ |
| A | 0557 | CD38 | 90 | Rat IgG2a, κ |
| A | 0558 | CD55 | RIKO-3 | Armenian<br>Hamster IgG |
| A | 0559 | CD63 | NVG-2 | Rat IgG2a, κ |
| A | 0560 | CD68 | FA-11 | Rat IgG2a |
| A | 0561 | CD79b | HM79-12 | Armenian<br>Hamster IgG |
| A | 0562 | CD83 | Michel-19 | Rat IgG1, κ |
| A | 0563 | CX3CR1 | SA011F11 | Mouse IgG2a,<br>κ |
| A | 0564 | Folate Receptor β | 10/FR2 | Rat IgG2a, κ |
| A | 0565 | MERTK | 2B10C42 | Rat IgG2a, κ |
| A | 0566 | CD301b | URA-1 | Rat IgG2a, λ |
| A | 0567 | Tim-4 | RMT4-54 | Rat IgG2a, κ |
| A | 0568 | XCR1 | ZET | Mouse IgG2b,<br>κ |
| A | 0570 | CD29 | HMβ1-1 | Armenian<br>Hamster IgG |
| A | 0571 | IgD | 11-26c.2a | Rat IgG2a, κ |
| A | 0573 | CD140a | APA5 | Rat IgG2a, κ |
| A | 0595 | CD11a | M17/4 | Rat IgG2a, κ |
| A | 0596 | ESAM | 1G8/ESAM | Rat IgG2a, κ |
| A | 0807 | CD200R | OX-110 | Rat IgG2a, κ |
| A | 0808 | CD193 | J073E5 | Rat IgG2a, κ |
| A | 0809 | CD200R3 | Ba13 | Rat IgG2a, κ |
| A | 0810 | CD138 | 281-2 | Rat IgG2a, κ |
| A | 0811 | CD317 | 927 | Rat IgG2b, κ |
| A | 0812 | CD105 | MJ7/18 | Rat IgG2a, κ |
| A | 0813 | CD9 | MZ3 | Rat IgG2a, κ |
| A | 0824 | P2X7R | 1F11 | Rat IgG2b, κ |
| A | 0825 | CD371 | 5D3/CLEC12A | Rat IgG2a, κ |
| A | 0827 | CD22 | OX-97 | Rat IgG1, κ |
| A | 0834 | CD39 | Duha59 | Rat IgG2a, κ |
| A | 0835 | CD314 | CX5 | Rat IgG1, κ |
| A | 0836 | DR3 | 4C12 | Armenian<br>Hamster IgG |
| A | 0837 | IL-33Rα | DIH9 | Rat IgG2a, $\kappa$ |
| A | 0839 | Ly49H | 3D10 | Mouse IgG1,<br>$\kappa$ |
| A | 0841 | Ly49D | 4E5 | Rat IgG2a, $\kappa$ |
| A | 0846 | CD185 | L138D7 | Rat IgG2b, $\kappa$ |
| A | 0848 | TIGIT | 1G9 | Mouse IgG1,<br>$\kappa$ |
| A | 0849 | CD80 | 16-10A1 | Armenian<br>Hamster IgG |
| A | 0850 | CD49a | HMα1 | Armenian<br>Hamster IgG |
| A | 0851 | CD1d | 1B1 | Rat IgG2b, $\kappa$ |
| A | 0852 | CD226 | 10E5 | Rat IgG2b, $\kappa$ |
| A | 0857 | CD34 | HM34 | Armenian<br>Hamster IgG |
| A | 0875 | TLR4 (CD284)/MD2<br>Complex | MTS510 | Rat IgG2a, $\kappa$ |
| A | 0876 | CD300c/d | TX52 | Rat IgG2b, $\kappa$ |
| A | 0877 | JAML | 4E10 | Armenian<br>Hamster IgG |
| A | 0881 | CD272 | 6A6 | Armenian<br>Hamster IgG |
| A | 0882 | PIR-A/B | 6C1 | Rat IgG1, $\kappa$ |
| A | 0883 | CD26 | H194-112 | Rat IgG2a, $\kappa$ |
| A | 0884 | DLL1 | HMD1-3 | Armenian<br>Hamster IgG |
| A | 0885 | CD270 | HMHV-1B18 | Armenian<br>Hamster IgG |
| A | 0890 | CD137L | TKS-1 | Rat IgG2a, $\kappa$ |
| A | 0891 | ENPP1 | YE1/19.1 | Rat IgG2b, $\kappa$ |
| A | 0892 | CD2 | RM2-5 | Rat IgG2b, $\lambda$ |
| A | 0895 | Mac-2 | M3/38 | Rat IgG2a, $\kappa$ |
| A | 0904 | CD31 | 390 | Rat IgG2a, $\kappa$ |
| A | 0905 | CD107a | 1D4B | Rat IgG2a, $\kappa$ |
| A | 0916 | CD124 | I015F8 | Rat IgG2b, $\kappa$ |
| A | 0917 | CD95 (Fas) | SA367H8 | Mouse IgG1,<br>$\kappa$ |

## References

1. Gherardi, M. M., Ramírez, J. C. & Esteban, M. IL-12 and IL-18 act in synergy to clear vaccinia virus infection: Involvement of innate and adaptive components of the immune system. Journal of General Virology vol. 84 1961–1972 (2003).

2. Pien, G. C., Satoskar, A. R., Takeda, K., Akira, S. & Biron, C. A. Cutting Edge: Selective IL-18 Requirements for Induction of Compartmental IFN-γ Responses During Viral Infection. J. Immunol. 165, 4787–4791 (2000).

3. Berghe, T. Vanden, et al. Simultaneous Targeting of IL-1 and IL-18 Is Required for Protection against Inflammatory and Septic Shock. Am. J. Respir. Crit. Care Med. 189, 282–291 (2014).

4. Mera, S. et al. Multiplex cytokine profiling in patients with sepsis. APMIS 119, 155–163 (2011).

5. Bourgon, N. et al. Cytokine Profiling of Amniotic Fluid from Congenital Cytomegalovirus Infection. Viruses 14, 2145 (2022).

6. Renneson, J. et al. IL-12 and type I IFN response of neonatal myeloid DC to human CMV infection. Eur. J. Immunol. 39, 2789–2799 (2009).

7. Akpan, U. S. & Pillarisetty, L. S. Congenital Cytomegalovirus Infection. StatPearls (2022).

8. Osterholm, E. A. & Schleiss, M. R. Impact of breast milk-acquired cytomegalovirus infection in premature infants: Pathogenesis, prevention, and clinical consequences? Rev. Med. Virol. 30, 1–11 (2020).

9. Namba, F. et al. Cytomegalovirus-related sepsis-like syndrome in very premature infants in Japan. Pediatr. Int. 64, e14994 (2022).

10. Kong, Y. et al. Sepsis-Induced Thymic Atrophy Is Associated with Defects in Early Lymphopoiesis. Stem Cells 34, 2902–2915 (2016).

11. Luo, M., Xu, L., Qian, Z. & Sun, X. Infection-Associated Thymic Atrophy. Front. Immunol. 12, 1947 (2021).

12. Ansari, A. R. & Liu, H. Acute Thymic Involution and Mechanisms for Recovery. Arch. Immunol. Ther. Exp. 2017 655 65, 401–420 (2017).

13. Lee, E. N. et al. Characterization of the expression of cytokeratins 5, 8, and 14 in mouse thymic epithelial cells during thymus regeneration following acute thymic involution. Anat. Cell Biol. 44, 14 (2011).

14. Pearse, G. Normal Structure, Function and Histology of the Thymus. http://dx.doi.org/10.1080/01926230600865549 34, 504–514 (2016).

15. Berthault, C. et al. Atrophy of primary lymphoid organs induced by Marek’s disease virus during early infection is associated with increased apoptosis, inhibition of cell proliferation and a severe B-lymphopenia. Vet. Res. 49, 1–18 (2018).

16. Ducimetière, L. et al. Conventional NK cells and tissue-resident ILC1s join forces to control liver metastasis. doi:10.1073/pnas.2026271118/-/DCSupplemental.

17. Shannon, J. P. et al. Group 1 innate lymphoid-cell-derived interferon-γ maintains anti-viral vigilance in the mucosal epithelium. Immunity 54, 276–290.e5 (2021).

18. Weizman, O. El, et al. ILC1 Confer Early Host Protection at Initial Sites of Viral Infection. Cell 171, 795–808.e12 (2017).

19. Sparano, C., et al. Embryonic and neonatal waves generate distinct populations of hepatic ILC1s. Sci. Immunol. 7, eabo6641 (2022).

20. Chen, Y. et al. Ly49E separates liver ILC1s into embryo-derived and postnatal subsets with different functions. J. Exp. Med. 219, (2022).

21. Lopes, N. et al. Tissue-specific transcriptional profiles and heterogeneity of natural killer cells and group 1 innate lymphoid cells. Cell reports. Med. 3, (2022).

22. Vosshenrich, C. A. J. J. et al. A thymic pathway of mouse natural killer cell development characterized by expression of GATA-3 and CD127. 7, 1217–1224 (2006).

23. S, G. et al. Murine thymic NK cells are distinct from ILC1s and have unique transcription factor requirements. Eur. J. Immunol. 47, 800–805 (2017).

24. Vargas, C. L., Poursine-Laurent, J., Yang, L. & Yokoyama, W. M. Development of thymic NK cells from double negative 1 thymocyte precursors. Blood 118, 3570–3578 (2011).

25. Cordes, M. et al. Single-cell immune profiling reveals thymus-seeding populations, T cell commitment, and multilineage development in the human thymus. Sci. Immunol. 7, (2022).

26. Schmitt, T. M., Ciofani, M., Petrie, H. T., Zúñiga-Pflücker, J. C. & Carlos Zúñiga-Pflücker, J. Maintenance of T Cell Specification and Differentiation Requires Recurrent Notch Receptor-Ligand Interactions. J. Exp. Med. J. Exp. Med 200, 469–479 (2004).

27. Lavaert, M. et al. Integrated scRNA-Seq Identifies Human Postnatal Thymus Seeding Progenitors and Regulatory Dynamics of Differentiating Immature Thymocytes. Immunity 52, 1088–1104.e6 (2020).

28. Stokic-Trtica, V., Diefenbach, A. & Klose, C. S. N. NK Cell Development in Times of Innate Lymphoid Cell Diversity. Front. Immunol. 11, 813 (2020).

29. Klose, C. S. N. et al. Differentiation of type 1 ILCs from a common progenitor to all helper-like innate lymphoid cell lineages. Cell 157, 340–356 (2014).

30. Gao, Y. et al. Tumor immunoevasion by the conversion of effector NK cells into type 1 innate lymphoid cells. Nat. Immunol. 18, 1004–1015 (2017).

31. Flommersfeld, S. et al. Fate mapping of single NK cells identifies a type 1 innate lymphoid-like lineage that bridges innate and adaptive recognition of viral infection. Immunity 54, 2288–2304.e7 (2021).

32. Park, E. et al. Toxoplasma gondii infection drives conversion of NK cells into ILC1-like cells. Elife 8, (2019).

33. Constantinides, M. G., McDonald, B. D., Verhoef, P. A. & Bendelac, A. A committed hemopoietic precursor to innate lymphoid cells. Nature 508, 397 (2014).

34. Daussy, C. et al. T-bet and Eomes instruct the development of two distinct natural killer cell lineages in the liver and in the bone marrow. J. Exp. Med. 211, 563–577 (2014).

35. Gordon, S. M. et al. The Transcription Factors T-bet and Eomes Control Key Checkpoints of Natural Killer Cell Maturation. Immunity 36, 55–67 (2012).

36. Friedrich, C. et al. Effector differentiation downstream of lineage commitment in ILC1s is driven by Hobit across tissues. Nat. Immunol. 2021 2210 22, 1256–1267 (2021).

37. Liu, B. et al. Severe influenza A(H1N1)pdm09 infection induces thymic atrophy through activating innate CD8+CD44hi T cells by upregulating IFN-γ. Cell Death Dis. 5, e1440–e1440 (2014).

38. Malaisé, M. et al. KLRG1+ NK cells protect T-bet-deficient mice from pulmonary metastatic colorectal carcinoma. J. Immunol. 192, 1954–1961 (2014).

39. Shin, S. B. & McNagny, K. M. ILC-You in the Thymus: A Fresh Look at Innate Lymphoid Cell Development. Front. Immunol. 12, 1605 (2021).

40. Lun, A. T. L., McCarthy, D. J. & Marioni, J. C. A step-by-step workflow for low-level analysis of single-cell RNA-seq data with Bioconductor. F1000R esearch 5, (2016).

41. Narni-Mancinelli, E. et al. Fate mapping analysis of lymphoid cells expressing the NKp46 cell surface receptor. Proc. Natl. Acad. Sci. U. S. A. 108, 18324–18329 (2011).

42. Ran, G. he, et al. Natural killer cell homing and trafficking in tissues and tumors: from biology to application. Signal Transduct. Target. Ther. 2022 71 7, 1–21 (2022).

43. Morillon, Y. M., Manzoor, F., Wang, B. & Tisch, R. Isolation and transplantation of different aged murine thymic grafts. J. Vis. Exp. 2015, (2015).

44. Liao, W., Lin, J. X., Wang, L., Li, P. & Leonard, W. J. Cytokine receptor modulation by interleukin-2 broadly regulates T helper cell lineage differentiation. Nat. Immunol. 12, 551 (2011).

45. Yoshimoto, T. et al. IL-12 up-regulates IL-18 receptor expression on T cells, Th1 cells, and B cells: Synergism with IL-18 for IFN-gamma production. J. Immunol. 161, 3400–3407 (1998).

46. Zhao, Y. et al. Biological Characteristics of Severe Combined Immunodeficient Mice Produced by CRISPR/Cas9-Mediated Rag2 and IL2rg Mutation. Front. Genet. 10, (2019).

47. Ferreira, A. C. F. et al. RORα is a critical checkpoint for T cell and ILC2 commitment in the embryonic thymus. Nat. Immunol. 2021 222 22, 166–178 (2021).

48. Qian, L. et al. Suppression of ILC2 differentiation from committed T cell precursors by E protein transcription factors. J. Exp. Med. 216, 884 (2019).

49. Woodfin, A., Voisin, M. B. & Nourshargh, S. PECAM-1: A Multi-Functional Molecule in Inflammation and Vascular Biology. Arterioscler. Thromb. Vasc. Biol. 27, 2514–2523 (2007).

50. Bigley, T. M. et al. Disruption of thymic central tolerance by infection with murine roseolovirus induces autoimmune gastritis. J. Exp. Med. 219, (2022).

## Online methods references

51. Tougaard, P. et al. TL1A Aggravates Cytokine-Induced Acute Gut Inflammation and Potentiates Infiltration of Intraepithelial Natural Killer Cells in Mice. Inflamm. Bowel Dis. 25, 510–523 (2019).

52. Kamimura, Y. & Lanier, L. L. Rapid and sequential quantitation of salivary gland-associated mouse cytomegalovirus in oral lavage. J. Virol. Methods 205, 53 (2014).

53. Li, W., Germain, R. N. & Gerner, M. Y. High-dimensional cell-level analysis of tissues with Ce3D multiplex volume imaging. Nat. Protoc. 2019 146 14, 1708–1733 (2019).

54. Hao, Y. et al. Integrated analysis of multimodal single-cell data. Cell 184, 3573–3587.e29 (2021).

55. Korsunsky, I. et al. Fast, sensitive and accurate integration of single-cell data with Harmony. Nat. Methods 2019 1612 16, 1289–1296 (2019).

